# Cryo-EM Structure of a Triazole α-Conotoxin GI Mimetic Bound to the Muscle-Type Nicotinic Acetylcholine Receptor

**DOI:** 10.64898/2026.08.31.748223

**Authors:** Oscar A. Shepperson, Michael J. Capper, Charlie Holdship, Oliver J. Melling, Nicola Wade, Michael A. Malone, Kirsty I.M. Arnott, Danielle C. Morgan, Thomas J. Piggot, Tiffany D. Morcom, Josephine A. Connah, Leonie M. Windeln, Christopher M. Timperley, Jeremy G. Frey, A. Christopher Green, Jesko Koehnke, Jonathan W. Essex, Andrew G. Jamieson

## Abstract

Disulfide-rich peptides possess exceptional potency and selectivity but are often limited by the instability and synthetic challenges associated with native disulfide bonds. Here, we report the design, synthesis, pharmacological evaluation, and structural characterisation of triazole-based peptidomimetics of the α-GI conotoxin, a selective antagonist of the muscle-type nicotinic acetylcholine receptor (nAChR). A series of 1,4- and 1,5-disubstituted triazole analogues were prepared entirely on resin using CuAAC and RuAAC chemistry to replace the native Cys3/13 disulfide bridge. Functional evaluation against human muscle nAChRs revealed that 1,5-triazole analogues retained low-nanomolar potency, with the lead mimetic exhibiting activity comparable to native α-GI. Cryo-electron microscopy of the lead compound bound to the muscle-type nAChR provided the first structure of a disulfide-isostere peptidomimetic in complex with a membrane receptor. The structure demonstrates that the 1,5-triazole reproduces the native peptide fold with high fidelity while contributing receptor-facing interactions not available to the native disulfide bridge. Molecular dynamics simulations further revealed conserved hydration networks and similar conformational sampling between the native peptide and lead mimetic. Together, these findings establish triazoles as effective disulfide surrogates and provide a structural framework for the rational design of stabilised conotoxin therapeutics.

## Introduction

Disulfide-rich peptides are among nature’s most potent and selective modulators of protein function, but their development is often limited by metabolic instability, synthetic complexity and the challenge of preserving bioactive conformations. The α-GI conotoxin, a disulfide-rich peptide derived from the venom of *Conus geographus*, is a potent and selective antagonist of the muscle-type nicotinic acetylcholine receptor (nAChR).^[1,2]^ A member of the α3/5 subfamily, α-GI is composed of 13 amino acids stabilised by two conserved disulfide bonds in a globular arrangement (I-III, II-IV).^[3]^ Unlike broader spectrum conotoxins, α-GI demonstrates minimal activity at neuronal nAChRs, making it a useful molecular probe for dissecting muscle receptor pharmacology for the development of subtype-specific inhibitors.^[4]^ Its well-defined pharmacological profile and high receptor specificity make it a valuable template for the development of therapeutic agents targeting neuromuscular disorders.^[5]^

Linear peptide therapeutics have historically encountered significant clinical limitations due to pharmacokinetic liabilities such as metabolic instability, poor membrane permeability, and susceptibility to proteolytic degradation. These challenges are particularly relevant for disulfide-rich peptide scaffolds, whose conformational integrity and biological activity often depend on labile disulfide bonds. As a result, considerable effort has been directed toward the development of peptidomimetics and disulfide isosteres capable of preserving native peptide topology while improving chemical and proteolytic stability. Beyond enhancing stability, structurally reinforced peptidomimetics may also provide opportunities to improve target engagement through additional or more favourable intermolecular interactions. Such approaches are especially relevant in the context of conotoxins and the nAChR, where subtle conformational and binding effects can strongly influence biological activity.^[6]^

Recent work by our group (2026) has elucidated the high-resolution cryo-electron microscopy (cryo-EM) structure of native α-GI bound to muscle-type nAChRs isolated from the pacific electric ray (*Tetronarce californica*).^[7]^ Intriguingly, an ambiguous cryo-EM desntiy was observed between the smaller disulfide bridge (I-III) of α-GI and the vicinal disulfide bond of the receptor C-loop. This observation was hypothesised to indicate a series of overlapped states in which intramolecular disulfide bonds have been broken and are forming intermolecular links. Under this model, neighboring disulfides within α-GI and the receptor C-loop can exchange connectivity, generating a transient covalent component to receptor engagement. In our previous studies investigating the systematic replacement of the disulfide bridges with disulfide isosteres, substitution of the Cys2-Cys7 disulfide bond in α-GI with a 1,5-triazole motif resulted in a marked reduction in conotoxin potency.^[6]^ In contrast, replacement of the Cys3/13 disulfide bridge with a 1,5-triazole revealed an equipotent peptidomimetic with increased stability.^[6]^ Relatively few studies examining disulfide bridge isosteres in the α-GI conotoxin have been reported. Previous work has explored lactam and thioether analogues with both studies reaching conclusions consistent with our own findings.^[8,9]^ In both cases a significant reduction in activity was observed upon replacement of the Cys2/7 disulfide, supporting its significance in α-GI conotoxin potency. The structural studies most relevant to the present work are those of Gray and co-workers (1996) and Craik and co-workers (1998) who elucidated the crystal and NMR structures of α-GI, respectively.^[3,10]^ In addition, triazole-based disulfide bridge replacements have been investigated in the conotoxin MrIA, where their effects on norepinephrine uptake inhibition were evaluated.^[11]^

To further develop our peptidomimetic chemistry and address the recent observations of disulfide cross-linking reported for the α-GI:nAChR cryo-EM structure, we sought to expand the chemical space around the 1,5-triazole as a peptidomimetic.^[6,7]^ Ultimately, we sought to determine if the key receptor interaction and binding mechanism of α-GI triazole peptidomimetics mimicked those for native α-GI.

Triazoles have long been key components of engineered peptides and small molecules, and have found wide applications across therapeutic areas including oncology, infectious diseases and protease inhibitors.^[12–14]^ Here, we report the complete on-resin synthesis of 1,4- and 1,5-disubstituted triazole-based peptidomimetics of the α-GI conotoxin, employing operationally straightforward click chemistry as a strategy to replace the native II-IV disulfide linkage. Specifically, we investigated triazole direction, leveraging the asymmetric nature of the triazole mimetic. Our synthetic analogues were evaluated for functional antagonism, resulting in the identification of a lead peptidomimetic. The structural basis for its interaction with *Tetronarce californica* nAChRs was elucidated through cryo-EM, providing structural insight into its mode of action and offering a platform for the future rational design of α-conotoxin-based therapeutics.^[7]^ This work represents the first reported cryo-EM structure of a peptidomimetic containing disulfide isosteres bound to a membrane receptor. Of particular interest we observe the maintenance of peptidomimetic conformation on binding as well as potential contribution to the interactions between the triazole moiety and the nAChR. This provides evidence for the unique capability of triazoles to not only effectively replace disulfide bridges but potentially modulate peptide-receptor interactions. Furthermore, structural and activity data further support a disulfide shuffling hypothesis through the design, synthesis and assessment of a small number of peptides functionalised with rationally designed covalent moieties.

## Results and Discussion

### Design and Synthesis of Native α-GI (1) and Conotoxin Peptidomimetics (2 – 8)

Through the synthesis of a series of α-GI conotoxin analogues, we sought to investigate two key hypotheses; (1) whether the asymmetry of the triazole influences its effectiveness as a disulfide mimic and, (2) whether chemically modified α-GI analogues could provide mechanistic insight into the proposed disulfide-shuffling process underlying conotoxin binding.

We initially focussed on expanding our understanding of the triazole mimetic structure-activity-relationship reported previously.^[6]^ We sought to leverage the regioisomeric and asymmetric nature of the triazole scaffold to probe its function as a disulfide bond mimic. We hypothesised that the retained potency of the previously reported 1,5-triazole analogue may result from favourable triazole–nAChR interactions and therefore investigated whether these interactions are affected by the orientation and regiochemistry of the triazole bridge.

The synthesis of globular conotoxin α-GI (**1**) was achieved using our previously reported methods and purified to yield α-GI (1) in moderate yields (7%) (**SI Table S2**).^[7]^

Peptidomimetics incorporating 1,4- or 1,5-disubstituted 1,2,3-triazole bridges in place of the II-IV disulfide bridge were synthesised by automated microwave-assisted Fmoc-SPPS. The commercial building blocks Fmoc-L-propargylglycine (Fmoc-Pra-OH) and Fmoc-L-azidohomoalanine (Fmoc-Aha-OH) were employed to provide side chains compatible for Copper or Ruthenium-catalysed Azide-Alkyne Cycloadditions (CuAAC/RuAAC) respectively. To facilitate on-resin disulfide bond formation Cys2/7 were protected with the orthogonally labile acetamidomethyl (Acm) protecting group. Following incorporation of the non-natural amino acids, the N-terminal Fmoc protected peptides were cyclised on-resin *via* the respective cycloadditions (**Scheme 1A/B**).^[15]^ These strategies selectively afforded on-resin 1,4- and 1,5-disubstituted 1,2,3-triazole-modified peptides. To enable concomitant on-resin removal of the acetamidomethyl (Acm) protecting group and disulfide bond formation between Cys2 and Cys7, *N*-chlorosuccinimide (NCS) was employed under mildly acidic conditions (**Scheme 1A/B**).^[16]^ The final desired triazole-containing peptides **2** – **5** were liberated from resin with global side-chain deprotection, and purified to yield the desired peptidomimetics in sufficient quantities for biological and structural studies (**2** – **5**: 2%) (**SI Table S2**).

Examination of the recently reported cryo-EM structure of α-GI bound to the nAChR led us to hypothesise that disulfide shuffling may contribute to the incredible potency of α-conotoxins.^[7]^ This is further supported by our previously reported observations of a complete loss of potency on replacement of the Cys2/7 disulfide bridge with a 1,5-triazole.^[6]^ We proposed that trapping the reaction intermediate through covalent functionalisation of a modified α-GI side chain could further test this hypothesis.^[17,18]^ Specifically, a modified side chain that, on binding, positions a reactive electrophile at an optimal distance from the proposed free thiol of Cys^193^ on the complementary loop of the nAChR α-subunits.

**Scheme 1.**
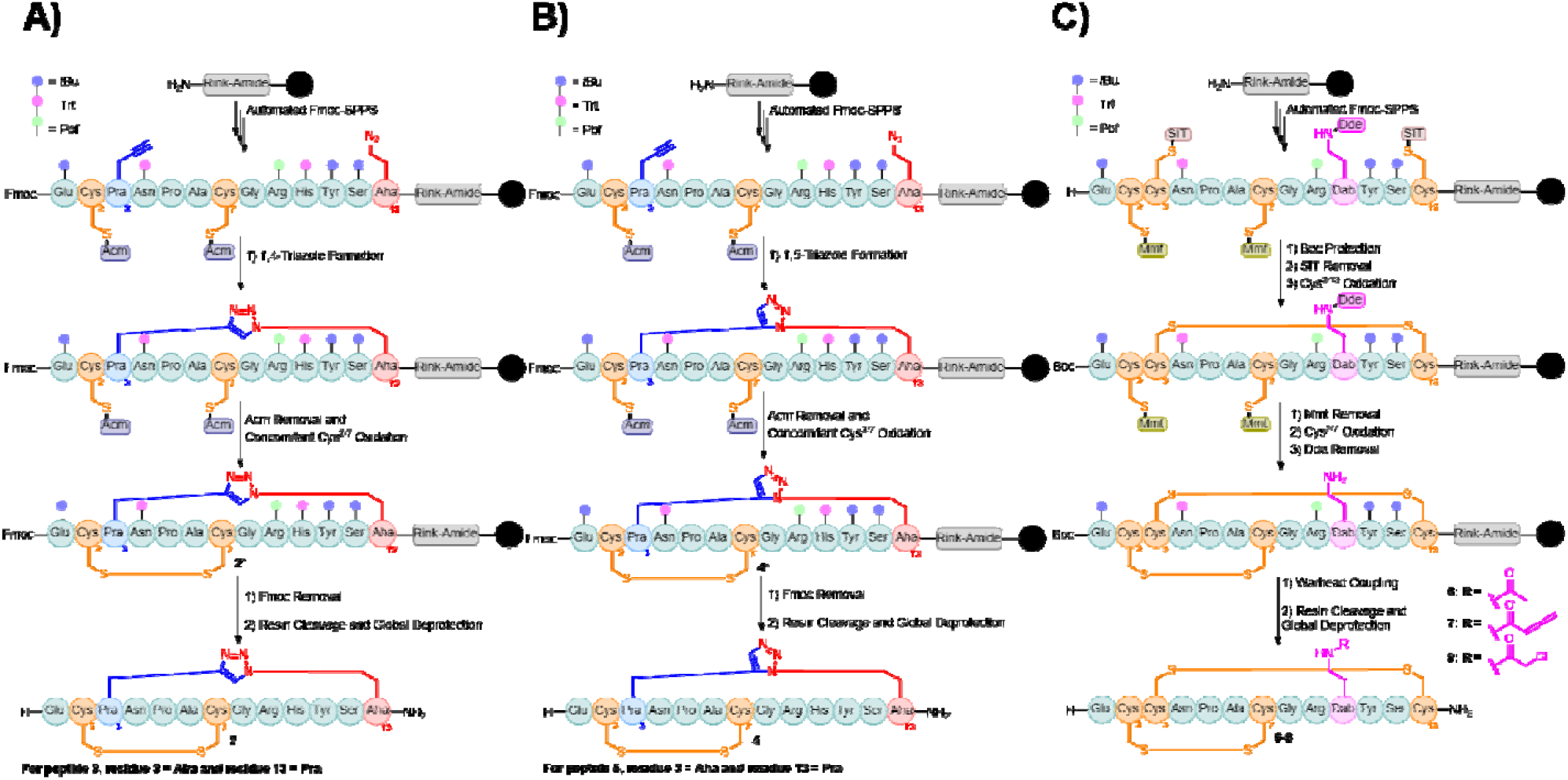
Entirely on-resin synthetic methodologies for the synthesis of **(A)** 1,4-triazole mimetics, **(B)** 1,5-triazole mimetics, and **(C)** covalently functionalised analogues. Cys and disulfide bridges (orange), Pra (blue), Aha (red), and Dab and covalent probes (pink). Detailed synthetic procedures can be found in the supplementary information (**SI Table S2**).

We employed the cryo-EM structure of native α-GI to determine the most well-suited residues to replace and orient molecular probes at a suitable distance. Through rational design the replacement of His10 with 2,3-diaminobutyric acid (Dab) was identified as the most suitable modification. Dab was positioned *via* alignment with the coulombic density observed for His10 in the cryo-EM structure of **1** (PDBID: 9SRL).^[7]^ Dab provided the additional advantage of being commercially available with orthogonal 1-(4,4’-dimethyl-2,6-dioxocyclohexylidene)-3-ethyl (Dde) protection. We envisaged Dde could be readily removed on-resin facilitating the functionalisation of the side chain amine with a small range of electrophilic functionalities.

Further rational design led to the selection of an amidated control moiety (**6**), alongside two probes, an allenamide (**7**) and chloroacetamide (**8**). These functionalities were identified as the most adequately placed (following attachment to the side chain of Dab10) for reactivity with the proposed free thiol of Cys193 of the complementary loop (**Figure 1**). Importantly, both of these moieties have been extensively used as covalent probes in a range of peptide and small molecule therapeutics.^[19,20]^ To enable incorporation of a thiol-sensitive reactive probe, a fully on-resin synthetic strategy for α-GI, compatible with Fmoc-SPPS, was required. Prior to incorporating Dab10 and our covalent probes we chose to standardise our on-resin methodology using α-GI (**1**). Our orthogonal synthesis of α-GI (**1**) employed Cys residues protected with 4-Methoxytrityl (Mmt) and *sec*-isoamyl mercaptan (SIT) at Cys2 and Cys7, and Cys3 and Cy**s**13 respectively (**Scheme 1C**).^[21–23]^ Following elongation of the linear peptide by automated Fmoc-SPPS, disulfide bridges were formed sequentially between Cys3 and Cys13, followed by Cys2 and Cys7. Following the removal of the orthogonal protecting groups of the respective Cys pairs, disulfide formation was facilitated by NCS.^[24]^ Key on-resin reactions were monitored by RP-HPLC and ESI-MS following cleavage of a small portion of the peptidyl resin (**SI Figures S1 – S7**). For the synthesis of analogues **6 – 8** linear peptides were synthesised employing our orthogonal strategy, with the N-terminal Fmoc protecting group replaced with Boc. Following completion of the on-resin orthogonal strategy, Dde was removed under nucleophilic conditions using hydrazine to reveal the free amine of Dab10.^[25]^ The covalent probe moieties, allenamide (**7, Figure 1A**) and chloroacetamide (**8, Figure 1B**), alongside an amide control (acetic acid) were conjugated to the Dab10 side-chain (**Scheme 1C**).^[19,20]^ Resin cleavage, global deprotection and subsequent purification yielded the three modified probe analogues **6, 7** and **8**, in analytically pure form (**6 – 8**: 2%) (**SI Table S2**).

**Figure 1.**
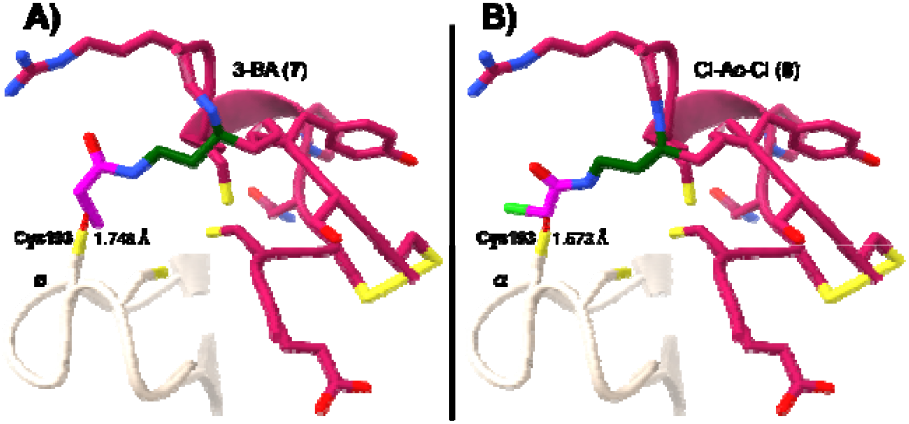
Rationally designed probe functionalities for α-GI analogues with **A)** allenamide (**7**) and **B)** chloroacetamide (**8**) functionalities built into cryo**-**EM structure of α-GI-nAChR (PDBID: 9SRL, white).^[7]^ Thiols and disulfide bridges (yellow), Dab (dark green), covalent probe (pink), halogen atoms (lime green) and intermolecular distance (red lines).

### Biological Evaluation of α-GI Analogues (1 – 8)

To determine the potency of the peptidomimetics synthesised herein, we chose to evaluate the *in vitro* activity against human receptors by assessing antagonism of the acetylcholine-induced, nAChR-mediated, increase of [Ca^2+^] in CN21γKO cells. CN21γKO cells expressing solely human adult muscle nAChRs are a well validated platform to assess the activity of nAChR antagonists.^[26]^ Addition of compounds **1 – 8** to CN21γKO cells produced concentration-dependent decreases in ACh-induced responses, indicative of nAChR antagonism (**Table 1**). Native α-GI (**1**) produced using our on-resin orthogonal strategy exhibited comparable potency to that of the α-GI produced *via* oxidative folding evidencing the desired isomeric canonical folding (**Table 1, Figure 2A**). Furthermore, the activity of native α-GI (**1**), and 1,5-triazole mimetic (**4**) aligned with previously reported data.^[6]^ Of particular interest, all triazole analogues (**2 – 5**) exhibited potent activities (**Table 1**). Triazole-containing peptides with the C-terminal Aha residues (**2** and **4**) exhibited greater potency than those with C-terminal Pra residues (**3** and **5**), with 1,5-triazoles (**4** and **5**) generally exhibiting greater potency than 1,4-triazoles (**2** and **3**) (**Table 1, Figure 2B**). This marginal difference in antagonism can be explained by the differential conformational flexibility of 1,4-vs. 1,5-triazoles as disulfide bridge mimetics.^[27]^ Our data further supports previous NMR data and *in silico* structural analysis establishing that the 1,5-triazole bridge is a more accurate mimetic of a disulfide bridge than the 1,4-triazole bridge, particularly in cases where the alkyne is positioned more N-terminally than the azide.^[28,29]^ This observation may arise from the 1,5-triazole bridge accurately reproducing the spatial arrangement of the native disulfide bond, such that the Cα-Cβ bond vectors of the Aha and Pra residues closely resemble those of the corresponding cysteine residues

**Table 1.** The calculated IC_50_ values from nAChR inhibition assays for peptides **1 – 8**. SEM refers to the standard error mean for at least four replicates.

| Compound | $IC_{50}$ (nM) ± SEM | $n_H$ |
| --- | --- | --- |
| **1 (folded)** | $3.41 \pm 1.00$ | -0.732 |
| **1 (orthogonal)** | $4.52 \pm 1.23$ | -0.792 |
| **2** | $8.00 \pm 1.06$ | -1.065 |
| **3** | $50.7 \pm 20.5$ | -0.967 |
| **4** | $4.24 \pm 1.09$ | -0.859 |
| **5** | $24.5 \pm 8.60$ | -1.100 |
| **6** | $18.9 \pm 4.10$ | -1.071 |
| **7** | $26.1 \pm 6.70$ | -1.003 |
| **8** | $2.78 \pm 1.06$ | -1.024 |

**Figure 2.**
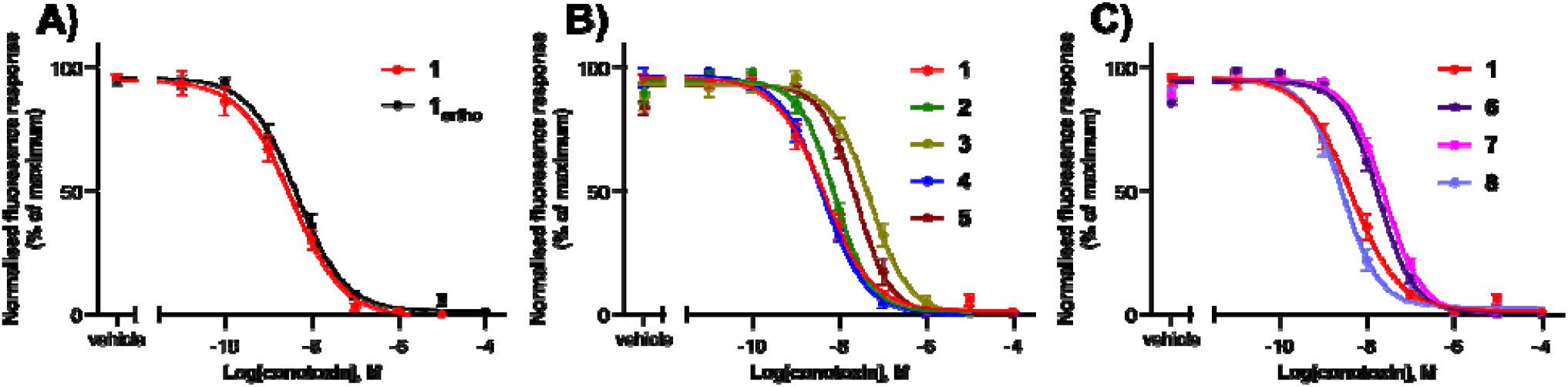
The acetylcholine-induced calcium response of CN21γKO cells pre-incubated with different concentrations of **A)** native α-GI, **B)** triazole mimetics, and **C)** covalent probes. Data are displayed as the mean ± standard error of the mean (SEM) of a minimum of four experiments performed in triplicate. Constrained sigmoidal dose-response curves, with a variable slope, were fitted for all compounds.

Analogues functionalised via the Dab side-chain **6 – 8** were investigated alongside the triazole mimetics for th**e**ir activity. Interestingly, the control analogue **6** exhibited a 5-fold reduction in potency (**Table 1**). However, the covalent pro**b**e functionalised analogues **7** and **8** produced variable activities. The analogue bearing a thia-Michael acceptor (**7**) exhibited diminished activity relative to the control, whereas the analogue bearing an S_N_2 probe (**8**) restored activity to the low-nanomolar range, with an IC_50_ comparable to native α-GI (**1**) (**Table 1, Figure 2C**). As these assays do not directly report covalent bond formation, the enhanced activity of **8** should be interpre**t**ed as functional support for productive positioning of the electrophile rather than direct evidence of covalent receptor modification.

We propose that these results, although indirect, support our hypothesis that disulfide shuffling may contribute to the exceptional potency of α-conotoxins. We hypothesise that peptide **7** was unable to restore activity to the same extent as peptide **8** owing to the reduced reactivity and increased rigidity associated with the sp-hybridised allenamide functionality.^[17, 30]^

### Structure of Mimetic 4 Bound to the Muscle-type nAChR

To determine the mechanism of binding of the triazole peptidomimetics and compare to that of native α-GI (**1**), we sought to resolve the cryo-EM structure of a bound peptidomimetic. Peptidomimetic **4** was selected for elucidation as it exhibited the greatest potency (IC50 = 4.24 ± 1.09 nM) of the synthesised peptidomimetics (**2 – 5**). Following the previously reported procedures, we determined the first high-resolution cryo-EM structure of an α-conotoxin peptidomimetic, **4**, bound to the muscle-type nAChR from *Torpedo californica*, a long-established model for nAChR structure and pharmacology, at 2.66 Å resolution (**Figure 3, SI Figures S8 and S9**).^[7]^

**Figure 3.**
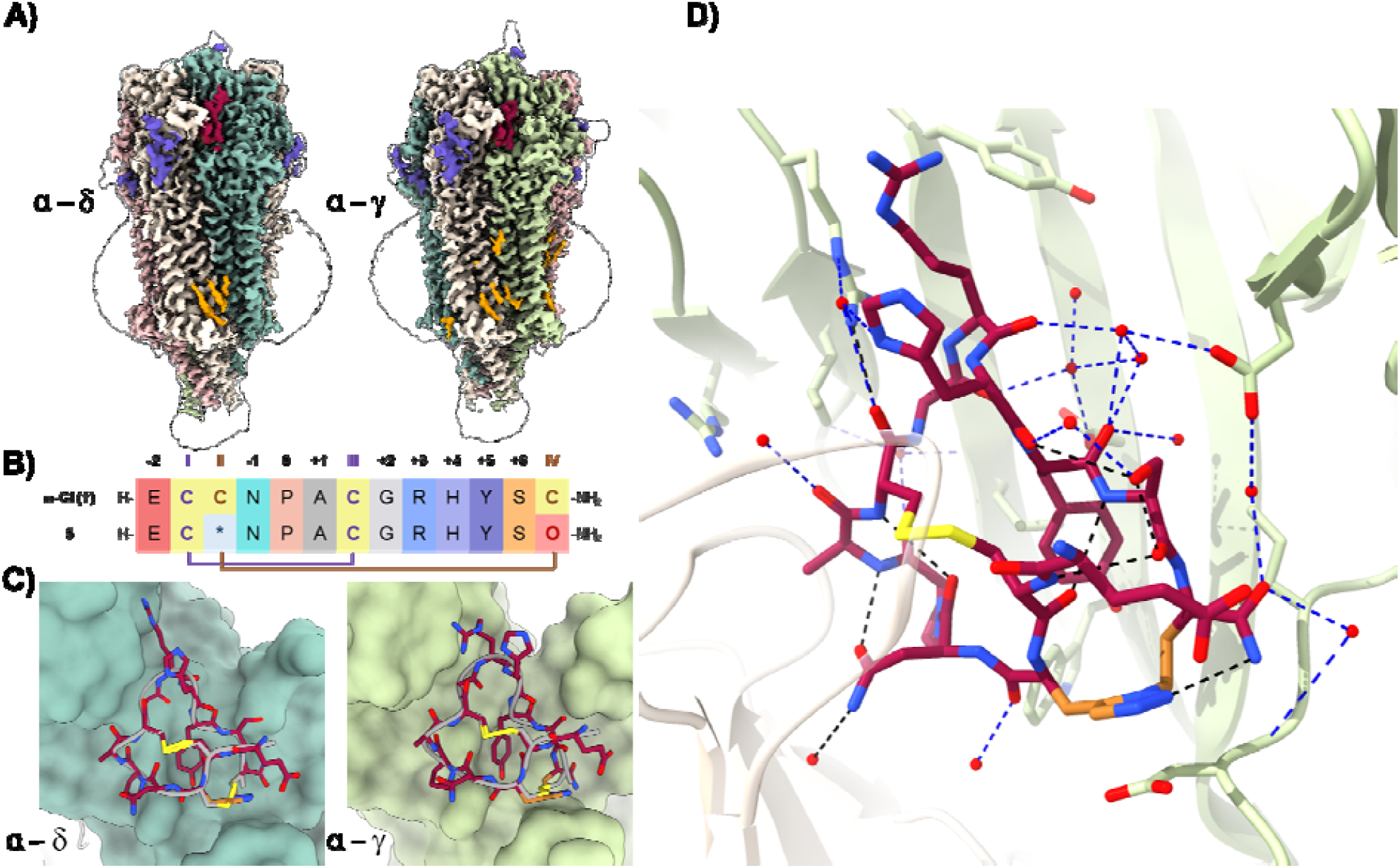
**A)** Unsharpened volumes of the nAChR bound to each **4** (crimson) at both the α – δ and α – γ binding sites (PDBID: 28WS), cholesterol (yellow) and glycosylations (purple). **B)** Sequence alignment of α-GI (**1**) with lead peptidomimetic **4. C)** Overlay of mimetic **4** with native conotoxin **1** (grey, PDBID: 9SRL) reveals overlapping structures possessing the same fold at the α – δ and α – γ sites respectively. **D)** Zoomed view of mimetic **4** bound to the nAChR highlighting the key hydrogen bonding interactions of mimetic **4** when bound to the α – γ site. 1,5-Triazole bridge (gold), water (red dots), disulfide bond (yellow) and hydrogen bonds (blue dashed lines). ^*^ = Propargylglycine and O = Azidohomoalanine.

### α-GI Peptidomimetic 4 Binding Interactions

The structure of the α-GI mimetic-bound nAChR allowed for the unambiguous placement of **4** at each of the two acetylcholine binding interfaces (**Figure 3**). Similarly to the interactions recently described by Capper *et al*. (2026), the binding of **4** precludes acetylcholine agonism, preventing pore opening and membrane depolarisation and results in flaccid paralysis due to inhibition of muscle contraction.^[7]^ To descri**b**e significant interactions, we have followed those rules defined by Capper *et al*. (2026) to assign the residues pertinent to the binding of **4**.^[7]^ Conotoxin mimetic **4** was divided at the cent**r**al Pro (defined as residue Pro-0). Non-cysteine residues toward the N-terminus have been assigned negative integers with residues toward the C-terminus assigned positive integers. Pro-0 also divides those residues interacting with the principal face (α) and complementary face respectively (δ or γ) (**Figure 3B**).

The subunit composition of the muscle-type nAChR gives rise to two distinct binding sites at the α – δ and α – γ interfaces. In a similar fashion to α-GI (**1**), Pro-0 of **4** forms a variety of hydrophobic interactions within the aromatic acetylcholine binding cage at the interface of the principal and complementary faces (α1Y93/α1W149 and δW57/δL121 or γW55/γL119).

Expanding upon the Pro-Ala lock, similar key interactions are observed for mimetic **4** as have been observed for nativ**e** α-GI (**1**). Namely, cationic Arg at the +3 position reaches upward toward the extracellular domain forming a salt bridge with D113 at the δ site and a cation-π interaction with Y111 at the γ site. Additionally, key interactions are present for Tyr at the +5 position. The side chain of Tyr+5 is ‘sandwiched’ in a hydrophobic pocket between δW57/δL121 and γW55/γL119 with the phenolic hydroxy forming weak hydrogen bonds with the side chains of δT38 and γT36. Finally, the solvent facing region of **4** is highly polar and the positively charged N- and C-termini maintain interactions with the highly charged glutamate region of the receptor. No significant interactions observed for the native conotoxin (**1**) are missing for the mimetic (**4**). Examining the water networks of the associated bound native (**1**) and mimetic (**4**) peptides reveals high homology, with consistent waters observed at both the α – δ and α – γ binding interfaces. The confidence of these waters was assessed *via* molecular dynamics (*vide infra*). Additional density is observed near the triazole and the complementary-face acidic residues δD180/γD174. Although the local resolution and chemical nature of this feature preclude definitive assignment, its position is consistent with a receptor-facing interaction involving the triazole-containing bridge. Closer evaluation of these interactions reveals them to be more prevalent at the α – δ interface than the α – γ interface, consistent with the previously reported variable binding affinities (**Figure 3C**).^[31]^ This observation is supported by the decreased distance measured between the triazole and δD180 (∼3.5 Å) when compared to γD174 (∼4.3 Å) (**SI Figure S10**).

Overlaying the peptide backbones of the peptides **1** and **4** reveals the 1,5-triazole mimetic as an exquisite disulfide bridge mimetic maintained upon receptor binding (**Figure 3D**). Overlaying the backbone Cα of **1** and **4** revealed non-equivalent root mean square deviations (RMSDs) at the α – δ and α – γ interfaces of 0.294 Å and 0.447 Å respectively. Specific inspection of the Cα and Cβ carbons of the bridge residues (residues 3 and 13) revealed a slightly larger RMSD, with values of 0.674 Å and 0.744 Å at the respective α – δ and α – γ (**SI Figure S11**). Although these values vary from those reported for the NMR structures of the native and 1,5 mimetics, they remain inside the experimental limits associated with assignment of the structures by cryo-EM.^[6,7]^

### Molecular Dynamics of Water Network for Native α-GI (1) and Lead Triazole Mimetic (4)

Analysis of the locations and occupancies of bound water molecules around the 1,5-triazole mimetic (**4**) was carried out by molecular dynamics (MD) simulations with grand canonical Monte Carlo (GCMC) enhanced sampling moves performed on the water molecules. An identical analysis was previously performed on hydration sites around the native α-GI conotoxin (**1**).^[7]^

At both binding sites, we see a high level of agreement in both the location and occupancy of the water molecules bound around the conotoxin between the native (**1**) and 1,5-triazole mimetic (**4**) α-GI conotoxins. At the α – δ binding site, the four highly conserved water molecules previously identified behind the WT conotoxin are all maintained when the mimetic is bound (**Figure 4A**). When the structures are aligned, on receptor backbone, the locations of these sites differ by at most 0.6 Å between the native (**1**) and the mimetic (**4**). The occupancies of the four sites are in good agreement, with the two cent**r**al hydration sites in the chain both occupied in 100% of simulations.

**Figure 4.**
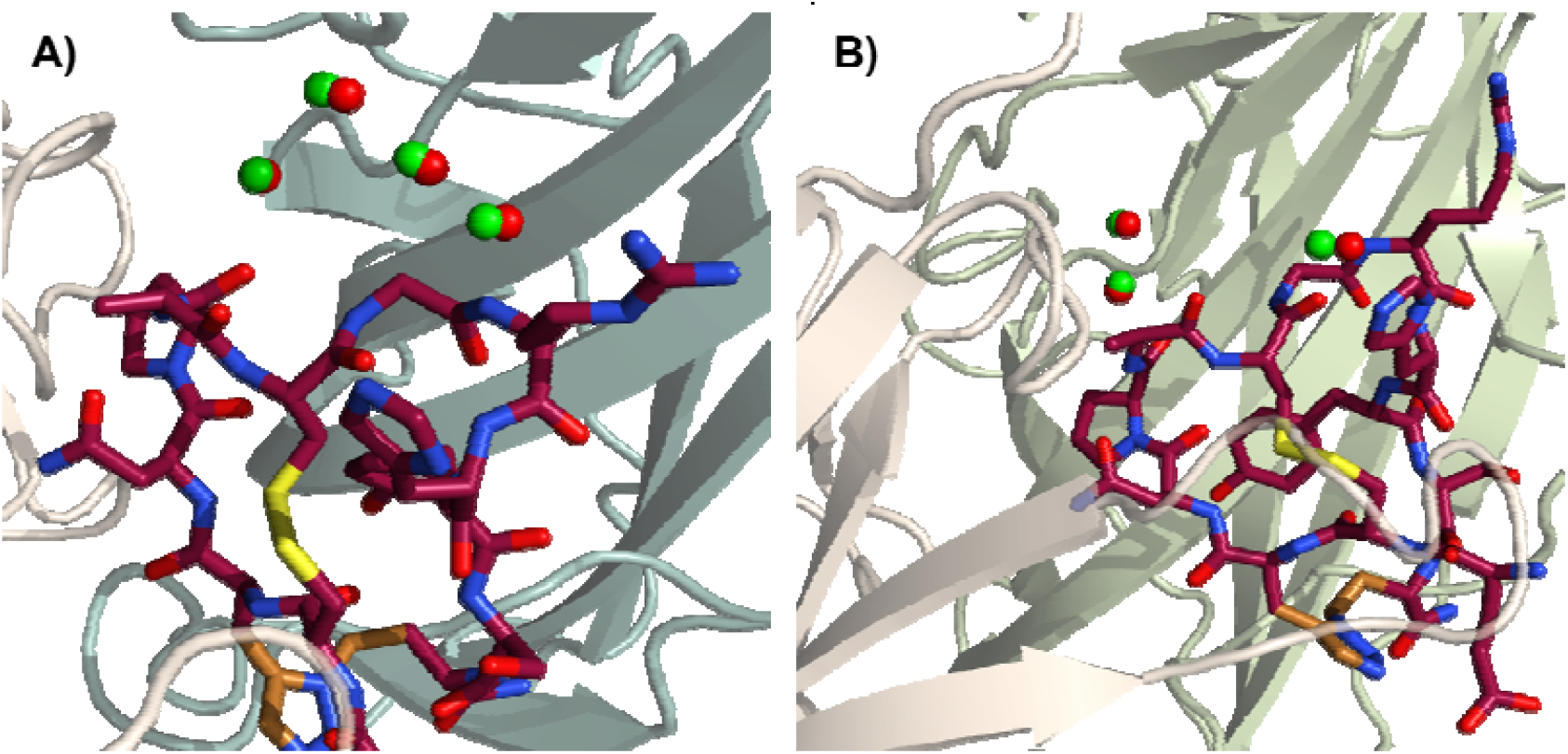
Comparison of hydration sites predicted by molecular dynamics simulations located behind the native (**1**) and 1,5-triazole (**4**) mimetic α-GI conotoxins bound at **A)** α – δ site, and **B)** α – γ site of the nAChR. Red and green spheres show the water locations around the native (**1**) and mimetic (**4**) conotoxins respectively.

At the α interface of the α – δ binding site, nine hydration sites were previously identified that were both within 5 Å of the native conotoxin and had an occupancy greater than 50%. When comparing to the hydration sites around the mimetic **(4**), four of the nine have a corresponding site within less than 1.0 Å, another four have a corresponding site between 1.0 Å and 2.0 Å away and one has a corresponding site 2.1 Å. We consider this to be reasonable agreement and not indicative of **t**he introduction of the triazole having any effect on the surrounding water network. At the δ interface, five of the seven hydration sites around **1** have a corresponding hydration site within 1.6 Å when the mimetic (**4**) is bound. As previously reported, the hydration pattern is typically noisier at this interface.^[7]^

For the α – γ binding site, similarly good agreement is seen.

Behind the conotoxin, three water locations are identified regardless of which conotoxin is bound (**Figure 4B**). When **t**he receptors are aligned each of these three sites sits within 1.0 Å of the corresponding site around the other conotoxin. Two of these waters have 100% occupancy in both simulations while the third has 60% occupancy in the native structure and 53% occupancy in the mimetic structure.

Around the rest of the α – γ binding site there were 10 waters identified when **1** was bound that had an occupancy of at least 50% and were within 5.0 Å of the conotoxin. Seven of these sites have a corresponding hydration site with at least 50% occupancy when mimetic **4** is bound and, all pairs of hydration sites are within 1.2 Å of each other. The lack of corresponding bound water locations around **4** for the other three sites can be ascribed to the threshold of 50% occupancy that we applied. Examination of water sites with occupancies below 50% reveals corresponding bound water molecules at the remaining three hydration sites. We conclude that replacing the disulfide bridge in native α-GI (**1**) with the 1,5-triazole mimetic in peptide (**4**) has no significant impact on the hydration pattern around the conotoxin in either of the two binding sites.

### Principal Component Analysis of α-GI and Mimetics 2 and 4

To investigate the proposed increased flexibility of the 1,4-mimetic compared to the 1,5-mimetic, we performed enhanced sampling molecular dynamics simulations (using REST2) in water of the wild type and both mimetics (compounds **1, 2** and **4**).^[32]^ We then employed principal component analysis (PCA) and k-means clustering to quantify the structural diversity of α-GI (**1**) and mimetics **2** and **4** in comparison with the nAChR bound conformations (PDBID: 9SRL), published α-GI NMR solution structure (PDBID: 1XGA), and published α-GI X-ray crystal structure (PDBID: 1NOT).

All analyses were performed using the backbone dihedral angles allowing for direct comparison between the wild type α-GI and the simulated mimetics. Plotting the projection of the trajectories of peptides **1, 2** and **4** onto the first two principal components of the wild type PCA, revealed similar as well as distinct distributions of conformations (**Figure 5**). Peptides **1** (wild type α-GI) and **4** (1,5-triazole mimetic) showed high similarity in terms of conformational sampling in the PCA plot. Of particular interest was that a proportion of the conformations of peptidomimetic **2** (1,4-triazole mimetic) occupied a unique region of conformational space. This indicated a population of unique conformations not observed for **1** and **4**.

**Figure 5.**
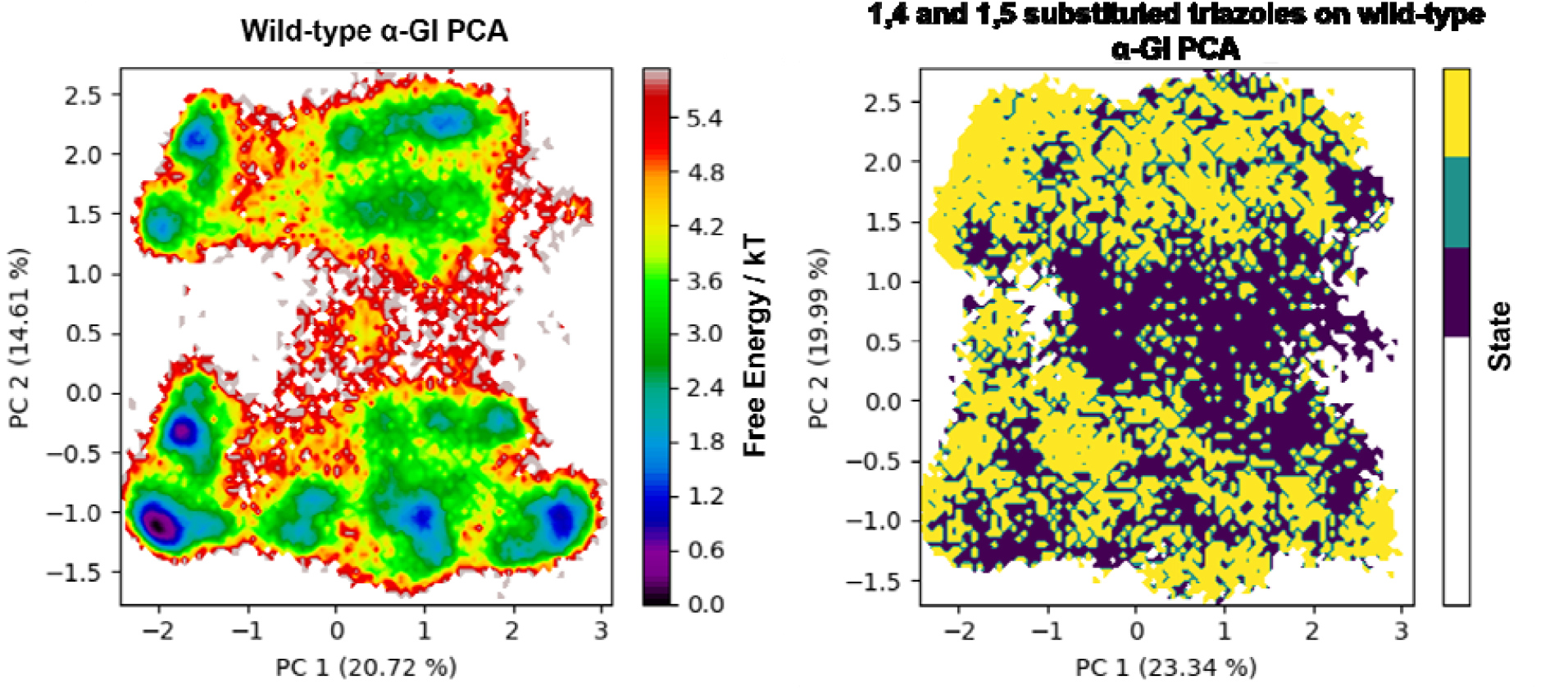
Comparison of principal component analysis plots of the wild type α-GI (**1**) (left), and the 1,4 and 1,5-triazole mimetics (**2** [purple] and **4** [yellow]) (right). All trajectories are plotted on the first two principal components of compound **1**. Overlapping regions indicate similar conformational distributions, with the 1-5 triazole mimetic (shown in yellow) having a greater overlap with the wild type indicating similar conformational distributions. The 1,4-triazole mimetic (shown in purple) has unique conformational distributions where PC2 ≈ 0.5 and PC1 is between -1 and 2 indicating a population of conformations not observed for compounds **1** and **4**. Shown in brackets is the percentage of variance in the trajectory that is captured by each of the principal components.

To further explore these conformations, we performe**d** k-means clustering on a combined trajectory of all three peptides. The resulting clusters were then split by their frequency in each simulation to see whether any were unique to each peptide. This revealed two clusters which were unique to mimetic **2** (clusters 4 and 6) and made up a combined 20% of the mimeti**c 2** structures, compared to only 2% of the α-GI structures and <1% of mimetic **4** structures (**SI Table S1**). Clusters 4 and 6 were also projected back on to the PCA plot to see which areas of **t**he conformational space they occupied and, cluster 4 overlapped with the unique portion of the distribution initially noted (**SI Figure S12**). On this basis, mimetic **2** is less similar i**n** its conformational distribution in solution than wild type α-GI (**1**) or mimetic **4**. This would be expected to result in a lower bin**d**ing affinity for mimetic **2** on both entropic and enthalpic grounds aligning with what is observed experimentally.

Each of the cluster centroids were compared to the known α-GI structures (compound **4** bound to nAChR, compound **1** bound to nAChR, and PDBIDs: 1NOT and 1XGA) by backbone dihedral angle RMSD to see into which clusters these structu**r**es would be placed, and how well our simulations match the known peptide conformational space. Interestingly, the cryo-EM of compounds **1, 4** and the crystal structure of **1** (1NOT) were all placed in the same cluster (cluster 3) while the NMR solution structure (1XGA), was put into cluster 1. Cluster 3 is the least well defined of the 11 clusters with lowest silhouette score (-0.11) (**SI Figure S13**) indicating a significant overlap with adjacent clusters, suggesting that there newly assign**e**d structures could be reasonably assigned to related cluste**r**s, including cluster 1. Wild type α-GI and both mimetics show significant conformation populations in the clusters to which the experimental structures have been assigned. On this basis, α-GI and the mimetics would all be expected to bind to nAChR but mimetic **2** sample additional conformations not populated by wild-type α-GI or mimetic **4**, consistent with its reduced potency relative to mimetic **4**.

## Conclusion

In this paper, we have developed a series of triazole-containing α-GI conotoxin peptidomimetics and established their structural and functional relationship to the native disulfide-rich peptide. Regioselective replacement of the Cys3/13 disulfide bridge with 1,2,3-triazoles revealed a clear dependence of biological activity on triazole orientation and substitution pattern, with 1,5-disubstituted analogues consistently outperforming their 1,4-disubstituted counterparts. The lead mimetic, **4**, retained low-nanomolar potency at the muscle-type nAChR, demonstrating adequate triazole positioning can effectively substitute for a structurally critical disulfide bridge. Cryo-EM analysis provided the first structure of a disulfide-isostere peptidomimetic bound to a membrane receptor and revealed an exceptional degree of structural conservation between native α-GI, **1**, and the peptidomimetic **4**. The mimetic preserved all key receptor contacts observed for the native peptide while introducing additional receptor-facing interactions associated with the triazole bridge. The consistency of water binding networks between α-G1 and the 1,5-triazole-containing peptidomimetic suggest that water mediated interactions are not contributing to differences in peptide binding affinity. Principal component analysis and clustering of enhanced sampling molecular dynamics simulations, indicate that the 1,4-triazole-mimetic **2** adopts additional solution-phase conformers to those of α-GI **1** and the 1,5-mimetic **4**. This reduction in preorganisation of the 1,4-triazole-containing peptidomimetic likely contributes to its lower binding affinity. Molecular dynamics simulations further demonstrated that the native hydration network and conformational landscape are largely maintained following disulfide replacement, providing a mechanistic explanation for the retention of biological activity. Additionally, the activity profiles of rationally designed covalent analogues provide functional support for a recently proposed disulfide-shuffling mechanism in α-GI:nAChR recognition. Although direct covalent bond formation remains to be definitively demonstrated, restoration of potency through strategic electrophile placement suggests that productive interactions with receptor thiols may contribute to the extraordinary affinity of α-conotoxins.

Collectively, these findings establish triazoles as more than passive disulfide surrogates, highlighting their capacity to preserve bioactive conformations while potentially contributing directly to receptor recognition. More broadly, this work provides a structural and mechanistic foundation for the rational design of stabilised disulfide-rich peptide therapeutics and chemical probes targeting ligand-gated ion channels.

## Supporting Information

The authors have cited additional references within the Supporting Information.^[6,15,16,22–24,26,33–61]^

## Data Code Availability

Cryo-EM maps and associated atomic models have been deposited in the EMDB and PDB respectively. α-GI mimetic bound nAChR (EMD-56926; PDBID: 28WS).

## Acknowledgements

We acknowledge the Scottish Centre for Macromolecular Imaging (SCMI) for access to cryo-EM instrumentation, funded by the MRC (MC_PC_17135, MC_UU_00034/7, MR/X011878/1) and SFC (H17007). We would like to thank Prof. David Beeson for supplying CN21□KO cells. We are grateful to Diamond for access and support of the cryo-EM facilities at the UK electron Bio-Imaging Centre (eBIC), proposal BI37630 and thank Prof. da Fonseca, Dr. E. Morris and Dr. K. Dent for their help in data collection. The authors acknowledge the use of the IRIDIS High Performance Computing Facility, and associated support services at the University of Southampton, in the completion of this work. All authors thank the Defense Threat Reduction Agency (DTRA, Research Programme Grant HDTRA12210001) for its financial support of this research, which was sponsored by the US Department of Defense. The content of the information does not necessarily reflect the position or the policy of the federal government, and no official endorsement should be inferred.

---

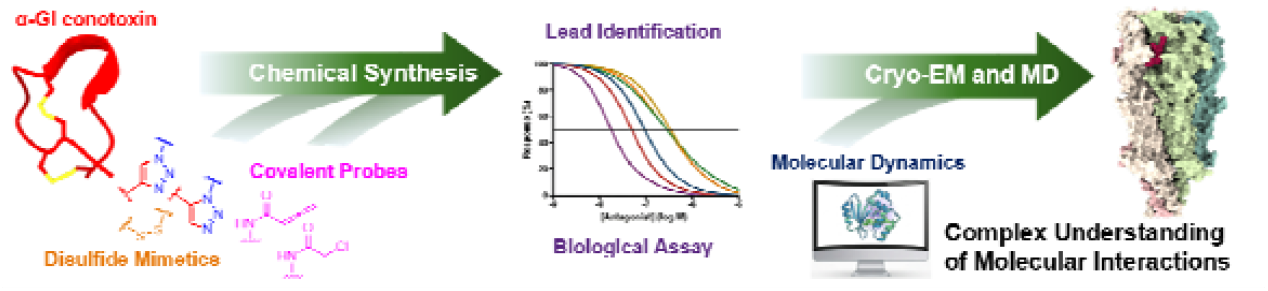

Triazole-based peptidomimetics of the α-GI conotoxin were developed as disulfide surrogates for the muscle-type nicotinic acetylcholine receptor (nAChR). A lead analogue retained native potency and enabled the first cryo-electron microscopy (cryo-EM) structure of a disulfide-isostere peptide bound to a membrane receptor. Alongside cryo-EM, molecular dynamics (MD) simulations were used to understand the structural basis of receptor recognition.

