## Supplemental Information for "Cryo-EM Structure of a Triazole α-Conotoxin GI Mimetic Bound to the Muscle-Type Nicotinic Acetylcholine Receptor"

[a]	Oscar A. Shepperson, Michael J. Capper, Nicola Wade, Michael A. Malone, Kirsty I. M. Arnott, Danielle C. Morgan, and Andrew G. Jamieson.
School of Chemistry, Advanced Research Centre, University of Glasgow, 11 Chapel Ln, Glasgow, G11 6EW, United Kingdom.

[b]	Charlie Holdship, Oliver J. Melling, Leonie M. Windeln, Jeremy G. Frey, and Jonathan W. Essex, School of Chemistry and Chemical Engineering, University of Southampton, Southampton SO17 1BJ, United Kingdom.

[c]	Thomas J. Piggot, Tiffany D. Morcom, Josephine A. Connah, Christopher M. Timperley, and A. Christopher Green, Defence Science and Technology Laboratory, Porton Down, Salisbury, SP4 0JQ, United Kingdom.

[d]	Jesko Koehnke, Institute of Food Chemistry, Leibniz University, Hanover, Germany

^+^ These authors contributed equally.

*.

### Supporting Information

### General Information and Methods

#### Reagents and Instrumentation

All reagents were purchased from commercial sources and used without further purification unless otherwise stated. Standard Fmoc-protected (Fmoc = 9-fluorenylmethyloxycarbonyl) amino acids were purchased from CEM Corporation and Pepceuticals, unless specifically stated differently below. Side chain protecting groups of *N^α^*-Fmoc amino acids were as follows; Fmoc-Asn(Trt)-OH (Trt = triphenylmethane), Fmoc-Arg(Pbf)-OH (Pbf = 2,2,4,6,7-pentamethyldihydrobenzofuran-5- sulfonyl), Fmoc-Cys(Trt), Fmoc-Glu(*t*Bu)-OH (*t*Bu = *tert*-butyl), Fmoc-His(Trt)-OH, Fmoc-Ser(*t*Bu)-OH, and Fmoc-Tyr(*t*Bu)-OH.

*N,N*-Dimethylformamide (DMF) and diethyl ether (Et_2_O) were purchased from Rathburn. Triisopropylsilane (TIPS), di-*tert*-butyl dicarbonate (Boc_2_O), hydrazine, chloroacetylchloride, chloro(cyclopentadienyl)bis(triphenylphosphine)ruthenium (CpRuCl(COD)), and ammonium bicarbonate (NH_4_HCO_3_) was purchased from Sigma-Aldrich, Merck. 2,2,2-Trifluoroethanol (TFE), ethane-1,2-dithiol (EDT), diisopropylcarbodiimde (DIC), ethyl 2-cyano-2-(hydroxyimino)acetate (Oxyma), Fmoc-Dab(Dde)-OH (Dde = 1-(4,4'-dimethyl-2,6-dioxocyclohexylidene)-3-ethyl), tris[(1-benzyl-1*H*-1,2,3-triazol-4-yl)methyl]amine (TBTA), Copper (I) iodide (CuI), Fmoc-Pra-OH, *N,N*-diisopropylethylamine (DIPEA), *N*-chlorosuccinimide (NCS), dithiothreitol (DTT), 3-butynoic acid (3-BA), 2,4,6-trimethylpyridine *sym*-collidine, *N*-ethoxycarbonyl-2-ethoxy-1,2-dihydroquinoline (EEDQ), sodium ascorbate (NaAsc), acetic anhydride (Ac_2_O), trifluoroacetic acid (TFA), and 2,6-lutidine were purchased from Fluorochem. Dichloromethane (CH_2_Cl_2_) was purchased from VWR. Acetonitrile (MeCN), piperidine, morpholine, and formic acid (FA) were purchased from Fisher Scientific. Fmoc-Aha-OH, Fmoc-Cys(SIT)-OH (SIT = *sec*-isoamyl mercaptan), Fmoc-Cys(Acm)-OH (Acm = acetamidomethyl), Fmoc-Cys(Mmt)-OH (Mmt = monomethoxytrityl) and aminomethyl Rink-Amide resin was purchased from Iris Biotech.

Analytical reverse-phase high-performance liquid chromatography (RP-HPLC) was performed on a Shimadzu RP-HPLC system with Shimadzu LC-20AT pumps, a Shimadzu SIL20A autosampler and a Shimadzu SPD-20A UV-vis detector using a Phenomenex Aeris Peptide XB-C18 (100Å, 5 µm, 150 × 4.6 mm). Compounds were eluted with linear gradients at column-dependent flow rates (1 mL/min for the Aeris), where buffer A = 0.1% TFA in H_2_O and buffer B = 0.1% TFA in MeCN. Data are reported as column retention time (t_R_) in minutes (mins). Crude peptides were purified by preparative RP-HPLC using an Agilent Technologies 1260 Infinity II Preparative LC System (monitoring at 214 nm and 280 nm), with a Phenomenex Gemini column 5 µm C18 column (100 Å, 250 x 21.2 mm). Peptides were eluted on linear gradients (10 mL/min) as determined by analytical RP-HPLC. The solvents employed were buffer; A = H_2_O + 0.1% TFA, and buffer; B = MeCN + 0.1% TFA.

Liquid chromatography-mass spectrometry (LCMS) was performed on a Thermo Scientific LCQ Fleet Ion Trap Mass Spectrometer using positive mode electrospray ionisation (ESI^+^), where buffer A = 0.1% TFA in 95% H_2_O/5% MeCN and buffer B = 0.1% TFA in 95% MeCN/5% H_2_O. A linear gradient of 5% – 95%B over 20 min with a flow rate of 1 mL/min was used with a Phenomenex Aeris Peptide XB-C18 (100 Å, 5 µm, 150 mm x 4.6 mm). Room temperature is abbreviated to r.t.

#### Peptide Synthesis

The abbreviation for molar equivalent is equiv. hereafter.

##### General Method 1A for Automated Fmoc-SPPS at Elevated Temperature

Peptides were synthesised batchwise as required on an Automated Biotage Initiator+ Alstra microwave synthesiser on a 0.1 mmol scale. The peptides were synthesised using Rink-Amide functionalised aminomethyl polystyrene (0.37 mmol/g loading, 0.270 g). Peptides were elongated in cycles of amino acid coupling followed by Fmoc removal. Standard Fmoc-protected amino acid (5 equiv., 0.2 M in DMF) coupling was achieved by treatment with DIC (5 equiv., 0.5 M in DMF) and Oxyma (5 equiv., 0.5 M in DMF) at 90 °C for 5 min. Cys residues were coupled at a reduced temperature (50 °C) to minimise racemisation, and Arg residues were double coupled.^[1]^ Fmoc removal was achieved by treatment with 20% morpholine + 5% formic acid in DMF (4 mL, *v/v/v*) at 90 °C for 2 min followed by 90 °C for 5 min. The resin was washed with DMF following Fmoc removals (4 x 4 mL), and after coupling (2 x 4 mL).

##### General Method 1B for Automated Fmoc-SPPS at Room Temperature

Peptides were synthesised batchwise as required on an Automated Biotage Initiator+ Alstra microwave synthesiser on a 0.1 mmol scale. The peptides were synthesised using Rink-Amide functionalised aminomethyl polystyrene (0.37 mmol/g loading, 0.270 g). Peptides were elongated in cycles of amino acid coupling followed by Fmoc removal. Standard Fmoc-protected amino acid (5 equiv., 0.2 M in DMF) coupling was achieved by treatment with DIC (5 equiv., 0.5 M in DMF) and Oxyma (5 equiv., 0.5 M in DMF) at room temperature for 60 mins. Arg residues were double coupled. Fmoc removal was achieved by treatment with 20% morpholine + 5% formic acid in DMF (4 mL, *v/v/v*) at room temperature for 1 x 5 min followed by 1 x 15 min with fresh reagents. The resin was washed with DMF following Fmoc removals (4 x 4 mL), and after coupling (2 x 4 mL).

##### General Method 2 for On-resin Removal of SIT from Cys by DTT

The resin bound peptide was treated with a solution of DTT (10 equiv.) in H_2_O/DIPEA/DMF (4:4:92, *v/v/v*, 5 mL, 12 x 10 min). Following the final treatment, as indicative with analysis by HPLC and ESI-MS following cleavage of a small portion of the resin, the resin bound peptide was thoroughly washed with DMF.^[2]^

##### General Method 3 for On-resin Disulfide Bond Formation by NCS

The resin bound peptide with exposed free thiols was treated with a solution of NCS (2 equiv.) in DMF (5 mL, 3 x 15 min). Following the final treatment, as indicative with analysis by HPLC and ESI-MS following cleavage of a small portion of the resin, the resin bound peptide was thoroughly washed with DMF.^[3]^

##### General Method 4 for On-resin Acid Mediated Removal of Mmt from Cys

The resin bound peptide was treated with a solution of CH_2_Cl_2_:TFA:TIPS (93:5:2, *v/v/v*, 5 mL, 15 x 5 min). Following the final treatment, as indicative by absence of any visible remaining orange colour of liberated Mmt protecting groups, the resin bound peptide was thoroughly washed with DMF.^[4]^

##### General Method 5 for N-terminal Fmoc to Boc Exchange

The *N^α^*-protected resin bound peptide was treated with a solution of 20% morpholine + 5% formic acid in DMF (2 x 5 mL, *v/v/v*) and thoroughly washed with DMF. The *N^α^*-unprotected resin bound peptide was treated with a solution of Boc_2_O (109.1 mg, 0.5 mmol, 10 equiv.) in DMF (10 mL), activated with DIPEA (174.8 µL, 1.0 mmol, 20 equiv.) for 2 h. Following treatment of the resin, with the reaction judged as complete by the confirmation of a negative result by a Kaiser test. The resin bound peptide was thoroughly washed with DMF.^[5]^

##### General Method 6 for On-resin 1,4-triazole Formation by Copper-catalysed Azide-Alkyne Cycloaddition (CuAAC)

The resin bound peptide (0.05 mmol, 1 equiv.) was swollen in DMF/TFE, 1:1 (*v/v*). To this, copper iodide (8.6 mg, 0.045 mmol, 0.9 equiv.), DIPEA (43.7 µL, 0.25 mmol, 5 equiv.), 2,6-lutidine (28.9 µL, 0.25 mmol, 5 equiv.), NaAsc (14.9 mg, 0.075 mmol, 1.5 equiv.) and TBTA (31.8 mg, 0.06 mmol, 1.2 equiv.) were added and the reaction heated by microwave (30 min, 55 °C). Upon reaction completion, the resin was washed with DMF, followed by CH_2_Cl_2_, and dried under vacuum.^[6]^

##### General Method 7 for On-resin Concomitant Removal of Acm from Cys and Oxidation by NCS

The resin bound peptide (0.05 mmol, 1 equiv.) was treated with a solution of NCS (14.7 mg, 0.11 mmol, 2.2 equiv.) in 2% TFA/CH_2_Cl_2_ (*v/v*) for 3 x 15 min. Following the final treatment, as indicative with analysis by HPLC and ESI-MS following cleavage of a small portion of the resin, the resin bound peptide was thoroughly washed with DMF, the N-terminal Fmoc removed, and thoroughly washed with DMF, followed by CH_2_Cl_2_, and dried under vacuum.^[7]^

##### General Method 8 for On-resin 1,5-triazole Formation by Ruthenium-catalysed Azide-Alkyne Cycloaddition (RuAAC)

The resin bound peptide (0.05 mmol, 1 equiv.), was swollen and degassed in a solution of DMF. To this CpRuCl(COD) (3.8 mg, 0.01 mmol, 0.2 equiv.) in DMF was added and the reaction heated by microwave (2 h, 70 °C). Upon reaction completion, the resin was washed with DMF, followed by CH_2_Cl_2_, and dried under vacuum.^[8]^

##### General Method 9 for On-resin Orthogonal Removal of Dde from Dab by Hydrazine

The resin bound peptide was treated with a solution of 2% hydrazine (*v/v*) in DMF (5 mL, 3 x 3 min). Following the final treatment, as indicative with analysis by HPLC and ESI-MS following cleavage of a small portion of the resin, the resin bound peptide was thoroughly washed with DMF.^[5]^

##### General Method 10A for Resin Cleavage and Global Deprotection Prior to Oxidative Folding

The resin-bound peptide was treated with a cleavage cocktail of TFA/H_2_O/TIPS/EDT (92/2/2/4, *v/v/v/v*, 10 mL) and agitated (2 h, r.t.). The cleavage solution was separated from the resin and its volume reduced under a flow of N_2_. Ice cold Et_2_O (15 mL) was used to precipitate the peptide. The precipitate was isolated by centrifugation, washed once more with ice cold Et_2_O, dissolved in MeCN/MQ H_2_O (2:8, *v/v*) and analysed by RP-HPLC and ESI-MS. Following determination of the desired peptidyl products, the solution was frozen and lyophilised.

##### General Method 10B for Resin Cleavage and Global Deprotection Following On-Resin Folding

The resin-bound peptide was treated with a cleavage cocktail of TFA/H_2_O/TIPS (95/2.5/2.5, *v/v/v*, 10 mL) and agitated (2 h, r.t.). The cleavage solution was separated from the resin and its volume reduced under a flow of N_2_. Ice cold Et_2_O (15 mL) was used to precipitate the peptide. The precipitate was isolated by centrifugation, washed once more with ice cold Et_2_O, dissolved in MeCN/MQ H_2_O (2:8, *v/v*) and analysed by RP-HPLC and ESI-MS. Following determination of the desired peptidyl products, the solution was frozen and lyophilised.

##### General Method 11 for Solution Phase Oxidation

The lyophilised peptide was solubilised in a solution of TFE/MQ H_2_O (1:1, *v/v*) buffered to pH ~9.5 by 0.1 M NH_4_HCO_3_. The respective linear conotoxins was added over the course of 2 h and left at r.t., with mixing, overnight, with complete oxidation confirmed as judged by analytical RP-HPLC and ESI-MS. Following complete oxidation, TFE was removed under reduced pressure, the solution acidified, frozen and lyophilised.^[9]^

#### Acetylcholine Calcium Response Assays Muscle-type nAChRs

Calcium response assays were carried out as previously established.^[8,10]^ CN21ɣKO cells expressing only the adult isoform of muscle-type nAChR were grown in Dulbecco’s Modified Eagle Medium (DMEM) supplemented with 10% Foetal Bovine Serum (FBS) and 1% penicillin/streptomycin in tissue culture treated flasks at 37 °C, 5% CO_2_. For the assay, cells were seeded to give a final density of 80,000 per well in black, clear bottom 96 well plates and allowed to attach and grow overnight. On the day of assay, media were removed and cells were loaded with calcium FLIPR 5 dye (Molecular Devices) made up in the proprietz buffer according to the manufacture’s instructions. This was then diluted in 1:1 with an equal mix of HBSS and DMEM buffered with 10 mM *N*-2-hydroxzethzlpiperiyine-*N*-2-ethanesulfonic acid (HEPES) pH 7.4. 1x HBSS Atropine was added to give a final concentration of 20 µM to prevent muscarinic receptor-mediated responses. Cells were then incubated at 37 °C for 25 min prior to the addition of ⍺-conotoxin GI or its mimetics, which were then incubated for a further 5 min. Plates were then placed in a FlexStation 3 multi-mode plate reader (Molecular Devices) and fluorescence (485 nm excitation, 525 nm emission) was measured over 60 s. Acetylcholine (150 µM), was added after 16 s and the resulting fluorescence upon intracellular calcium ion concentration increase was measured. The average of the 15 s baseline was subtracted from the maximum fluorescence and this background corrected peak fluorescence plotted against dose to form the concentration response curve from which an IC_50_ could be calculated using GraphPad Prism 10 and a four-parameter logistic fit. Concentrations were run in quadruplicate on each plate, and each experiment was run at least three times independently.

#### Purification of *T. californica* nAChR

*T. californica* nAChR was purified following established protocols.^[11,12]^ Briefly, frozen electric organ tissue (Medix Biochemica) was thawed before being resuspended in 20 mM NaH_2_PO_4_ pH 7.4, 400 mM NaCl, and 0.5 mg/mL *N*-ethyl maleimide and homogenised in a blender. Membranes were isolated by centrifugation at 200,000 g for 25 min, resuspended in 20 mM Tris pH 11, 80 mM NaCl, 1 mM EDTA, 20% sucrose (*w/v*) and allowed to incubate for 30 min on ice. Membranes were subsequently harvested and homogenised in 20 mM NaH_2_PO_4_ pH 7.4, 100 mM NaCl three times undergoing multiple washes. Membranes were snap frozen in liquid nitrogen and stored at -80 °C.

Membranes were resuspended in 20 mM NaH_2_PO_4_ pH 7.4, 80 mM NaCl and solubilised in 1.5% Triton X-100 (*v/v*) for 1 h before centrifugation at 200,000 g for 30 mins to remove insoluble material. Solubilised material was incubated with immobilised ligand affinity resin (ATM-sepharose) for 1 h before washing in 20 mM Tris pH 8, 100 mM NaCl, 1 mM EDTA, 0.05% DDM:CHS (*w/v*). Protein was eluted in the same buffer that contained 100 mM carbachol and 10 mM β-mercaptoethanol. Protein containing-fractions were concentrated and desalted to remove carbachol and β-mercaptoethanol.

Protein samples for cryo-EM were then incubated with 0.1% LMNG:CHS (*w/v*) on ice for 1 h before size-exclusion chromatography (SEC) on a Superose 6 10/300 (Cytiva) in 20 mM HEPES pH 7.5, 100 mM NaCl, 0.1 mM EDTA, 0.00075% (*w/v*) LMNG, 0.00015% CHS (*w/v*) and 0.00015% GDN (*w/v*).

#### Cryo-EM Grid Preparation and Data Acquisition

SEC purified nAChR was concentrated to 5-10 µM and kept on ice. 1,5-triazole ⍺-Conotoxin GI was dissolved to 10 mM in SEC buffer before being added to purified receptor at 30-fold excess and incubated on ice for 30 min. The complex was then concentrated to *ca.* 6 mg/mL (20-30 µM) prior to grid preparation and 0.3 mM FFC8 was added to minimise preferential particle orientation. 3 µL of protein was added to Quantifoil® 1.2/1.3 300 Cu mesh grids that had been glow discharged in air for 30 s at 35 mA. Samples were plunge-frozen in liquid ethane using a Vitrobot with blot force 3 for 3 s at 8 °C and 95% relative humidity. Frozen grids were stored in liquid nitrogen before screening and data collection.

Grids were screened on a JEOL F200 at SCMI (Glasgow, UK) and those demonstrating thin ice were subsequently taken forward for high-resolution data collection on an eBIC KriosIV (Diamond, Oxford, UK). Data collection statistics are summarised in **Tables S3 and S4**.

#### Data Processing and Model Refinement

Data were processed in cryoSPARC v4.6.2 and individual workflows are shown in more detail in **Figures S8 – S9**.^[13]^ Each dataset was processed in a similar manner beginning with Patch Motion Correction and Patch CTF estimation. Particles were picked with templates generated from screened grids on a JEOL F200 and extracted with 4x binning prior to two rounds of 2D classification. Junk particles were discarded, and *ab initio* models were produced from both the initially discarded junk particles to act as sinks and the particles following classification. Heterorefinement was performed on all particles in the 2D classes and those fitting the sole remaining good class were re-extracted with 2x binning. Particles were CTF refined and subjected to a masked 3D classification to remove the detergent micelle which yielded multiple maps. The best group of particles were then extracted un-binned following reference-based motion correction and subjected to non-uniform refinement yielding a final volume which was then sharpened.

The apo *T. californica* nAChR model^[14]^ (PDB:7SMM) was fit into the map and refined using Phenix^[15]^ followed by manual building and the addition of the conotoxins, lipids, glycosylations and waters in Coot^[16]^ then by further real-space refinement in Phenix. Final models were deposited in the PDB and validated. Protein interface area was calculated using ePISA.^[17]^ Pore analysis was carried out using MOLEonline.^[18]^ Images were created in ChimeraX.^[19]^ Interactions were analysed in Arpeggio.^[20]^

#### Molecular Dynamics for Water Simulations of Compounds (1 and 4)

The computational water placement was performed using grand canonical Monte Carlo (GCMC) simulation using both instantaneous and nonequilibrium move proposals as implemented in the grand Python library.^[21,22]^ To set up the systems, all glycans were removed from the cryo-EM structure and the subunits were cut such that only the extracellular region was simulated. For the α subunits, the proteins were cut from residue Pro211 onwards, the β subunit was cut from residue Leu218 onwards, the δ subunit from residue Leu226 onwards and the γ subunit from residue Leu220 onwards. In each case, an *N*-methyl amide cap was added to ensure the termini remained neutral. The system was protonated at pH 7.4, solvated and parameterized in accordance with the AMBER 14ffSB forcefield for the protein and ions and the TIP3P forcefield for the water molecules.^[23,24]^ Sodium and chloride ions were added to both ensure the system was neutral and achieve an ion concentration of 0.15 M.

The simulations were performed in OpenMM 8.0.0.^[25]^ Equilibration was performed over a series of steps combining both GCMC moves and molecular dynamics (MD) simulation. An initial 10,000 GCMC moves were followed by 100 cycles of 1,000 GCMC moves and 10 fs MD in the NVT (number volume temperature) ensemble. 5 ns of MD in the NPT (number pressure temperature) ensemble was then used to ensure the system volume was equilibrated correctly. Finally, 500 cycles of 1 ps NVT simulation and 200 GCMC moves completed the equilibration. The production simulations were performed using nonequilibrium GCMC (GCNCMC) moves – each simulation consisting of approximately 15,000 moves with 5 ps MD sampling between moves. To analyse the water placement within the simulations, the trajectories were decorrelated such that every 10^th^ frame was retained and a hierarchal average-linkage clustering algorithm, as implemented in SciPy, was used to determine the locations and occupancies of each cluster.^[26]^ The GCMC moves were only performed within a spherical region within each system. For each binding site, the sphere is centred on the midpoint of the Cα atoms of two selected residues. In all cases the sphere radius was 14 Å.

The GCMC sphere was defined as the midpoint between the Cα atoms of residues Asn4 and Tyr11.

Harmonic positional restraints were used to prevent movement of the α-GI conotoxin in the α – δ binding site, as was observed in initial simulations. These restraints were applied to all Cα atoms in both the α-conotoxin and the nAChR receptor and a force constant of 100 kJ nm^-2^ was used. No restraints were applied to the α – γ binding site.

#### Molecular Dynamics for Principal Component Analysis (PCA) of Compounds 1, 2 and 4

Each of the α-GI wild type, 1,4 and 1,5-triazole mimetics (compounds **1**, **2** and **4** respectively) were simulated by enhanced sampling molecular dynamics using the REST2 implementation. Simulations were performed using Gromacs (version 2021.2) patched with Plumed (version 2.7.4) and with the CHARMM36m forcefield.^[27–29]^ All simulations were seeded from the first solution NMR structure of α-GI (1XGA) and the H++ web server was used to protonate the structure to physiological pH (7.4).^[30]^

The mimetic structures were created from the protonated solution NMR structure with the II/IV disulfide replaced by a triazole group using the builder tool in PyMOL.^[31]^ To make the structure compatible with pdb2gmx, new residues were added to the CHARMM forcefield. Where applicable, standard CHARMM forcefield parameters were used for the newly added residues, while additional dihedrals were fitted to quantum mechanical data using the FFTK plugin of VMD.^[32]^

All structures were solvated using TIPS3P water and sodium and chloride ions were added to neutralize the charges and achieve an ionic concentration of 0.15 M.^[33]^ The structures were then energy minimized by 100,000 steps of steepest descent followed by 5,000 steps of conjugate gradient descent. Systems were subsequently equilibrated for 100 ps in the NVT ensemble using a modified Berendsen thermostat followed by an additional 100 ps in the NPT ensemble using the Parinello-Rahman barostat. The production REST2 simulations were run in the NVT ensemble using the Nose-Hoover thermostat. These consisted of 13 replicas which were geometrically spaced with effective temperatures ranging between 310-900 K. Each replica was run for 1 μs using a 2 fs timestep, with exchange attempts between replicas being made every 100 simulation steps. Coordinates were written out every 10 ps and only the trajectory of the base replica was used for analysis.

#### Clustering Analysis

Clustering was performed using the k-means NANI algorithm on the cosines of the backbone dihedral angles.^[34]^ Using only the backbone dihedral angles allowed for comparison between the wild type α-GI and the triazole mimetics. The number of cluster centres to use was chosen by analysing the silhouette scores and Davies-Bouldin Indices (DBI) of the wild type α-GI simulation clustered using a range of cluster numbers from 4 to 20. The resulting elbow plot (**Figure S3**) indicated that 11 cluster centres was optimal for this system. For consistency between simulations, 11 cluster centres were therefore used for all subsequent analyses of α-GI and its mimetics.

##### Silhouette Score

Silhouette score is a metric quantifying the quality of a given cluster by averaging the silhouette coefficients of each point in the cluster.^[35]^ This silhouette coefficient is given by:

$$s= \frac{b-a}{max(a,b)}$$

Where $a$ is the mean distance between a point and all other points in its cluster and $b$ is the mean distance between that point and all other points in the next nearest cluster. Ranging from -1 to 1, high positive values indicate good agreement between a point and the rest of its cluster. Scores around 0 suggest no clear separation between adjacent clusters and negative values indicate that a datapoint has a negative impact upon its cluster cohesion and separation.

##### Davies-Bouldin Index (DBI)

The Davies-Bouldin Index (DBI) is a global measure of cluster similarity and gives the average similarity between each cluster and its most similar cluster.^[36]^ Given by:

$$DBI= \frac{1}{k}\sum_{i=1}^{k} \max_{i\neq j} R_{ij}$$

Where k is the number of clusters, $R_{ij}$ is the similarity measure for two clusters i and j, itself given by:

$$R_{ij}= \frac{s_{i}+s_{j}}{d_{ij}}$$

With $s_{i}$ and $s_{j}$ being the cluster diameters and $d_{ij}$ the distance between the two cluster centroids. By averaging over the maximum cluster similarities, the DBI gives a good indication of the separation between each of the clusters with values closer to zero meaning better separation.

### Supporting Information Schemes, Tables and Figures

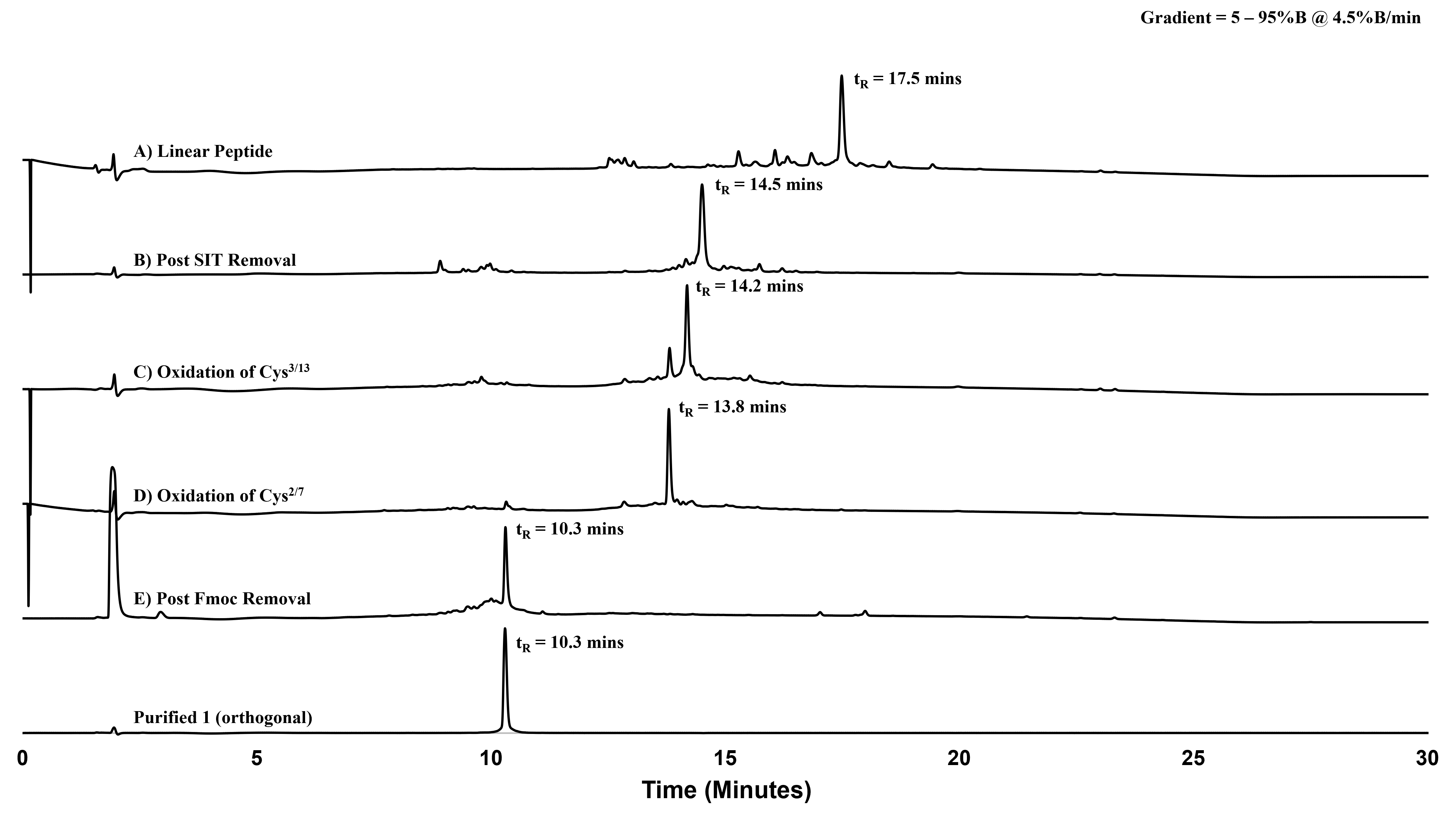

**Supporting Information Figure S1.** RP-HPLC traces (214 nm) following reaction progression for the orthogonal synthesis of **1**. Phenomenex Aeris Peptide XB-C18 (100 Å, 5 µm, 150 mm x 4.6 mm), linear gradient 5% – 95%B over 20 min (*ca.* 4.5%B/min) at 1 mL/min.

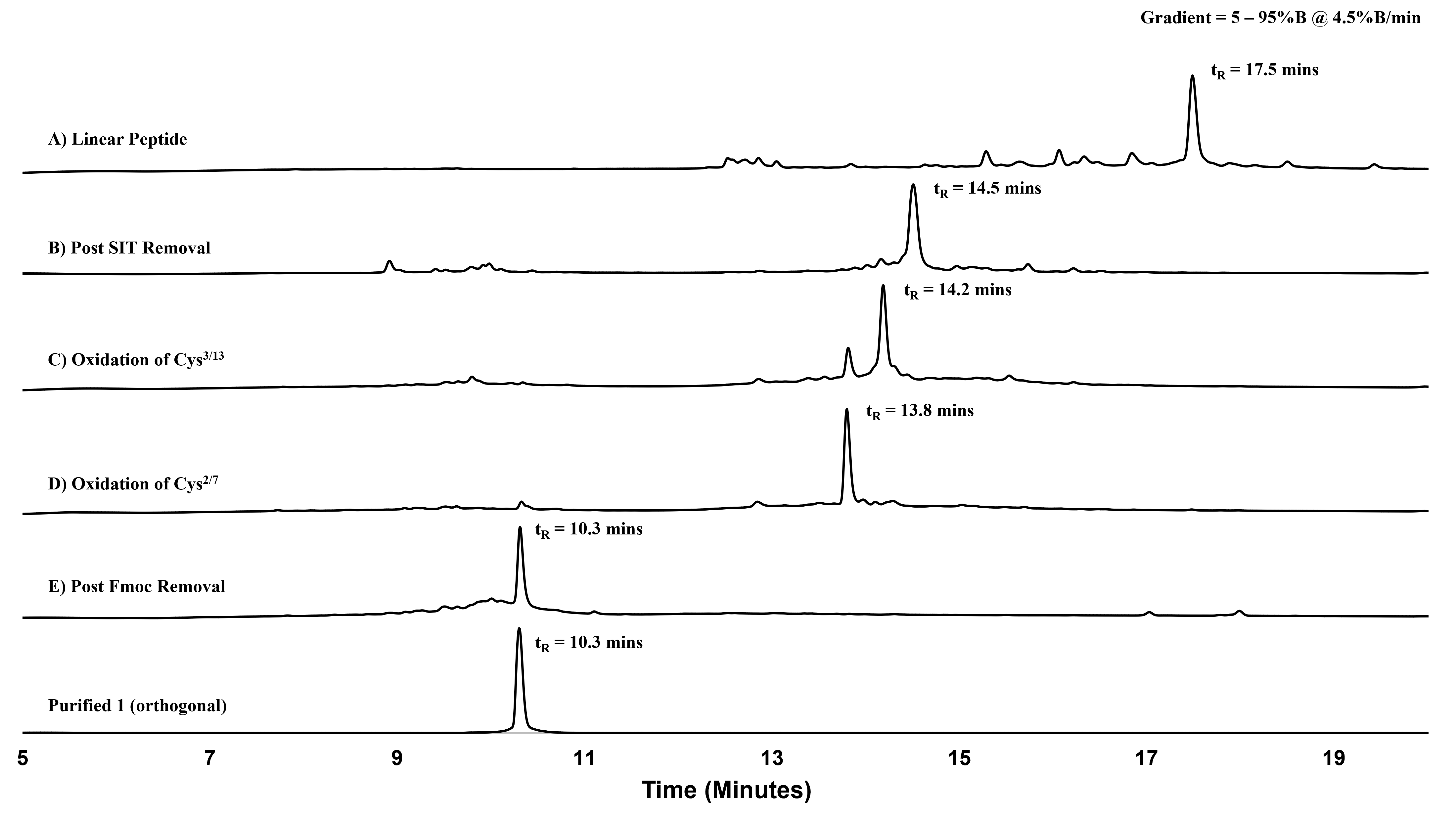

**Supporting Information Figure S2.** Zoomed RP-HPLC traces (214 nm) following reaction progression for the orthogonal synthesis of **1**. Phenomenex Aeris Peptide XB-C18 (100 Å, 5 µm, 150 mm x 4.6 mm), linear gradient 5% – 95%B over 20 min (*ca.* 4.5%B/min) at 1 mL/min.

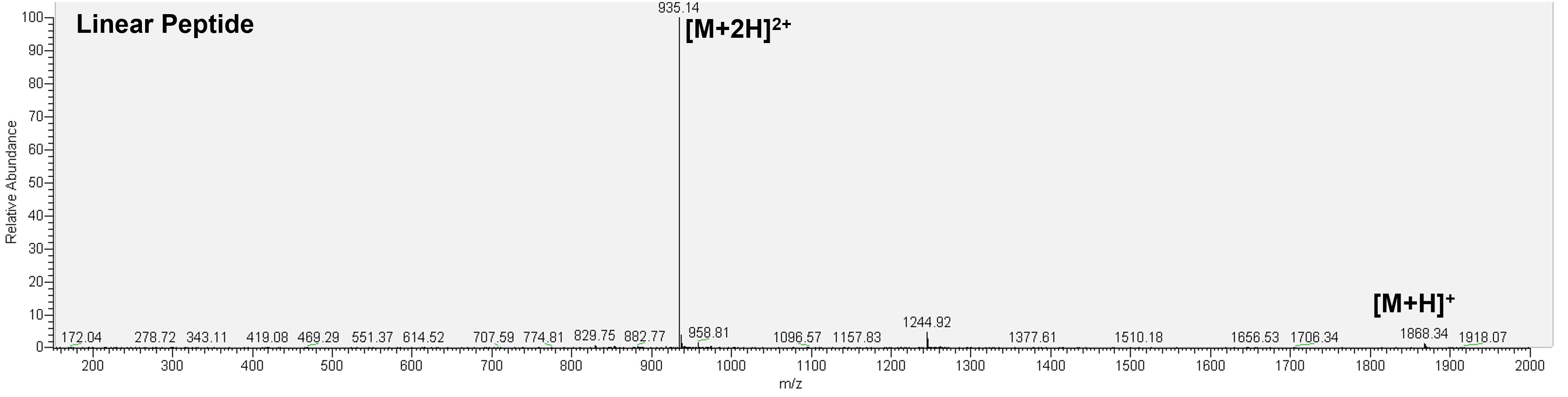

**Supporting Information Figure S3.** ESI-MS of major peptide intermediate product of RP-HPLC trace **SXA (Linear Peptide)**, mass calculated for [C_80_H_114_N_20_O_20_S_6_ + H] 1868.27; deconvoluted mass observed: 1867.81 ± 0.66. Charge states; 935.14 [M+2H]^2+^, 1868.34 [M+H]^+^.

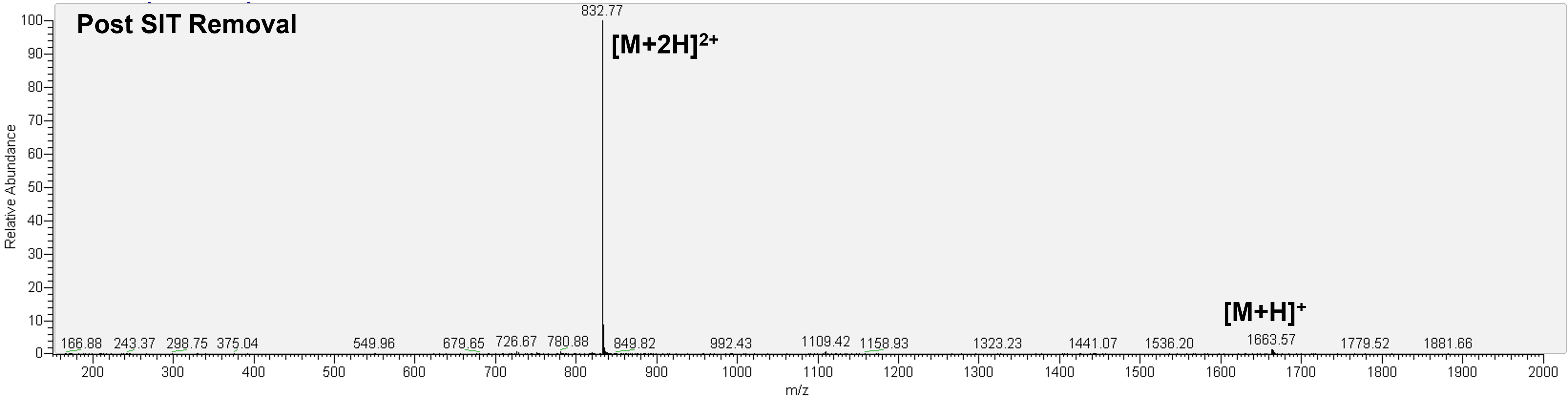

**Supporting Information Figure S4.** ESI-MS of major peptide intermediate product of RP-HPLC trace **SXB (Post SIT Removal)**, mass calculated for [C_70_H_94_N_20_O_20_S_4_ + H] 1663.88; deconvoluted mass observed: 1663.06 ± 0.69. Charge states; 832.77 [M+2H]^2+^, 1663.57 [M+H]^+^.

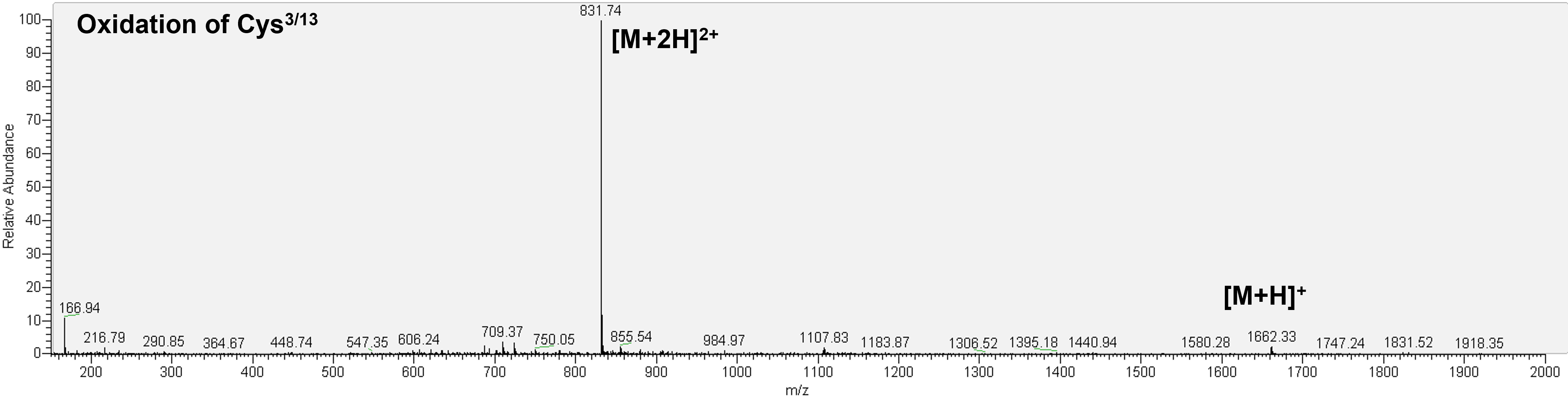
**Supporting Information Figure S5.** ESI-MS of major peptide intermediate product of RP-HPLC trace **SXC (Oxidation of Cys^3/13^)**, mass calculated for [C_70_H_92_N_20_O_20_S_4_ + H] 1661.87; deconvoluted mass observed: 1661.41 ± 0.41. Charge states; 831.74 [M+2H]^2+^, 1662.33 [M+H]^+^.

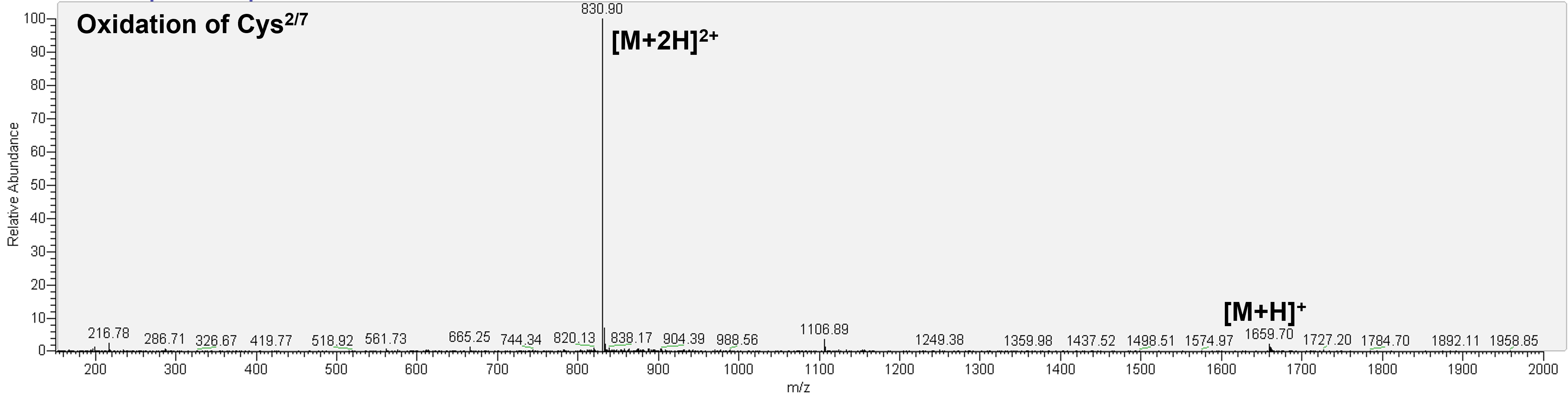

**Supporting Information Figure S6.** ESI-MS of major peptide intermediate product of RP-HPLC trace **SXD (Oxidation of Cys^2/7^)**, mass calculated for [C_70_H_90_N_20_O_20_S_4_ + H] 1659.85; deconvoluted mass observed: 1659.25 ± 0.78. Charge states; 830.90 [M+2H]^2+^, 1659.70 [M+H]^+^.

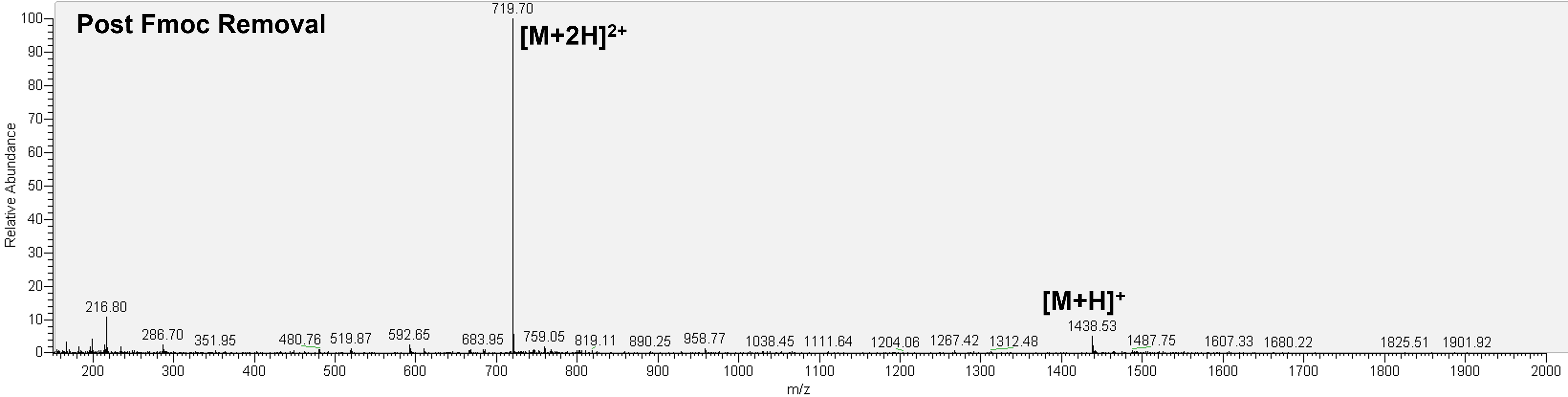

**Supporting Information Figure S7.** ESI-MS of major peptide intermediate product of RP-HPLC trace **SXE (Post Fmoc Removal)**, mass calculated for [C_55_H_80_N_20_O_18_S_4_ + H] 1437.61; deconvoluted mass observed: 1437.47 ± 0.09. Charge states; 719.70 [M+2H]^2+^, 1438.53 [M+H]^+^.

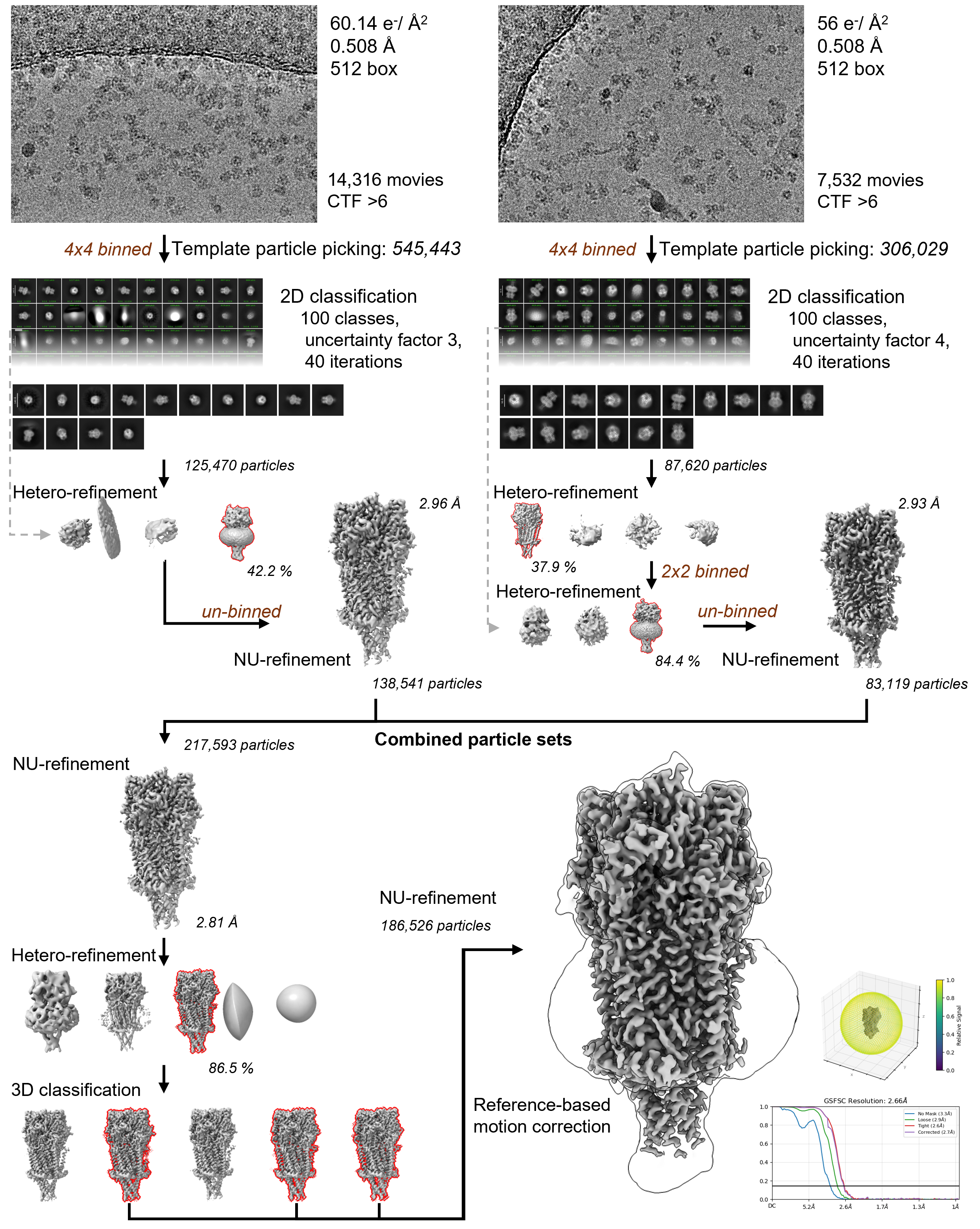

**Supporting Information Figure S8.** Cryo-EM data processing flowchart with orientation statistics and FSC curve for the final α-GI:triazole (**4**) bound nAChR structure.

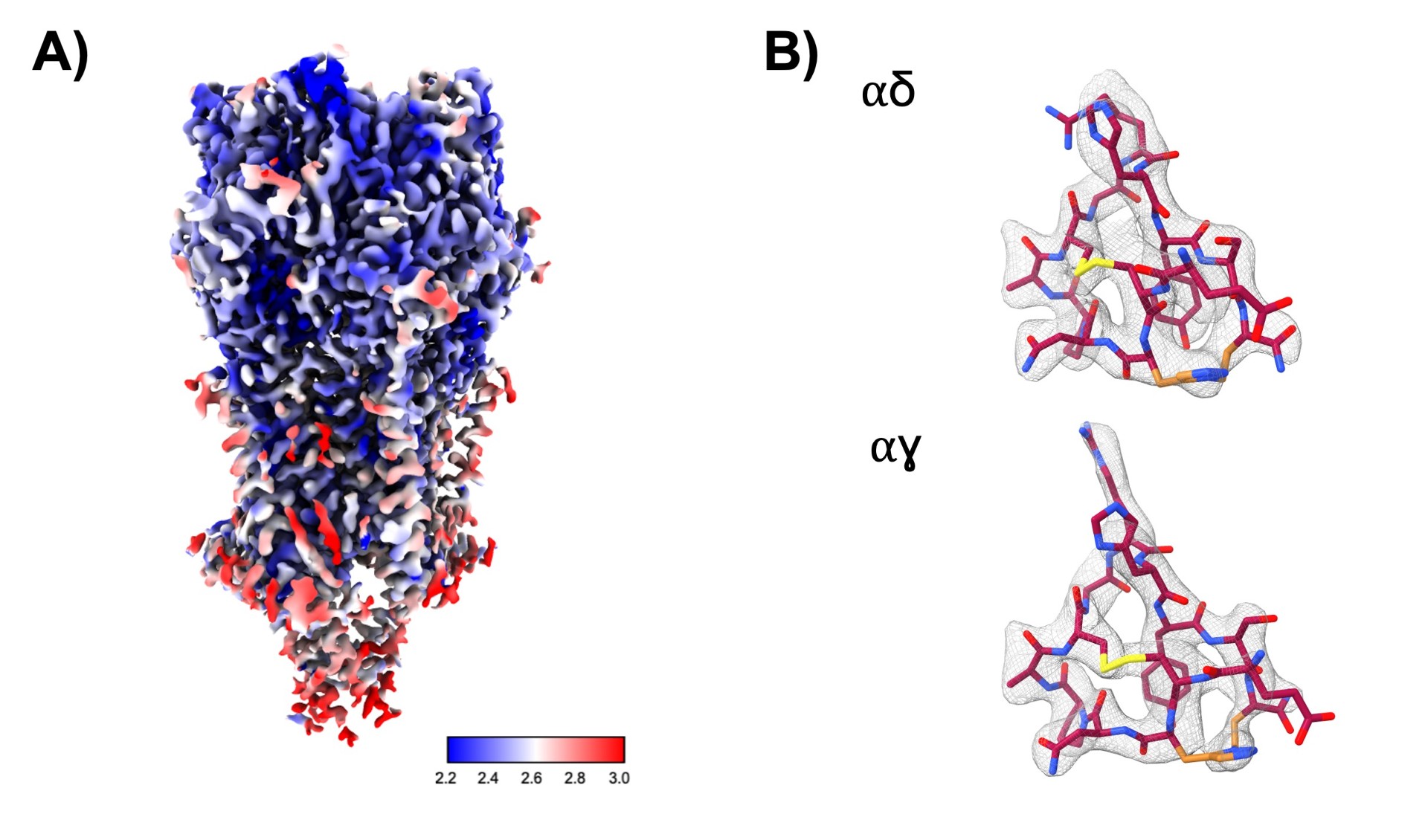

**Supporting Information Figure S9. A)** Local resolution maps for α-GI:triazole (**4**) bound nAChR structure. **B)** Sharpened cryo-EM density map contoured and surrounding α-GI:triazole (**4**) at αδ and αγ sites respectively (upper and lower) (PDBID: 28WS).

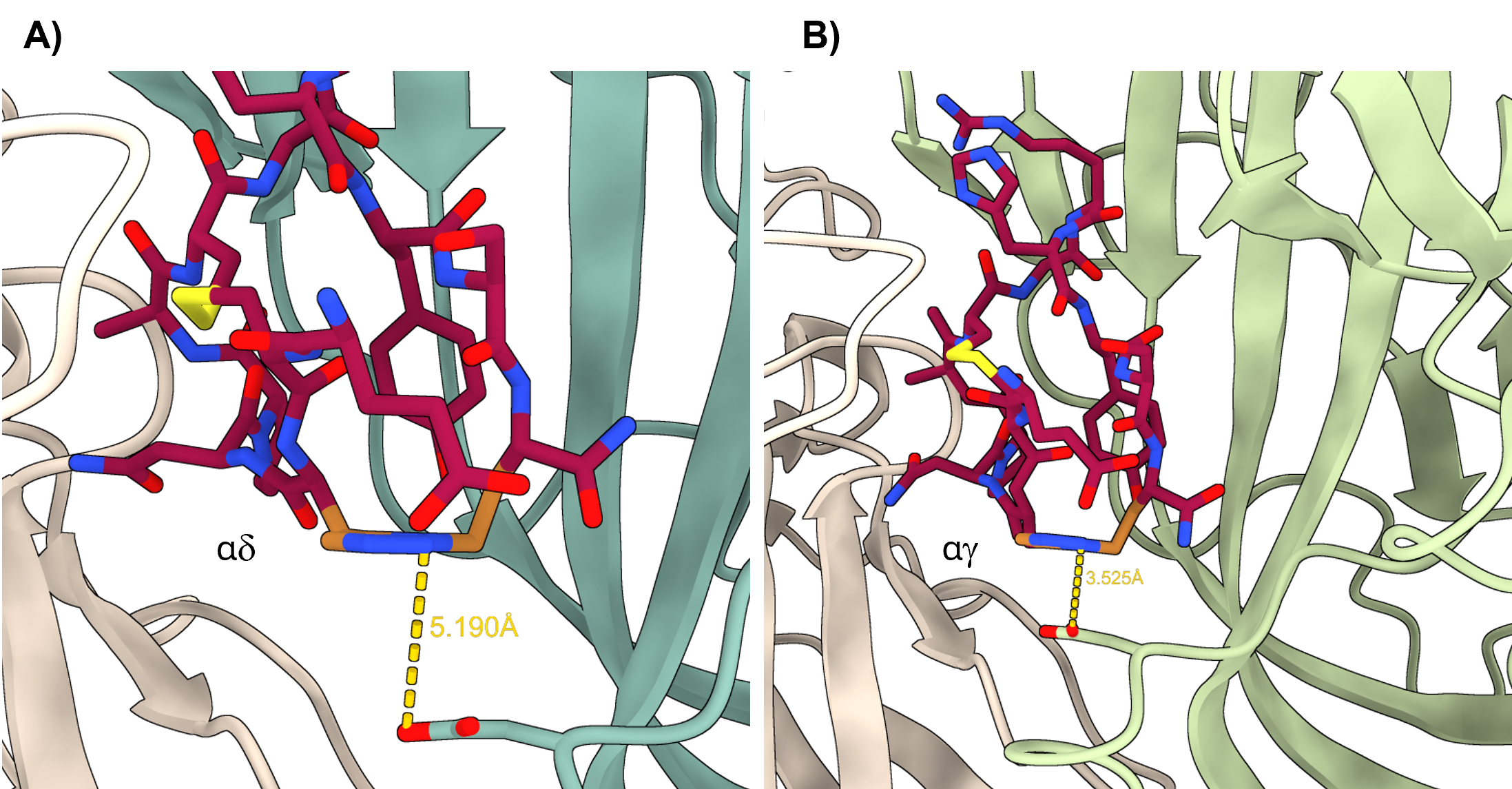

**Supporting Information Figure S10.** Interaction of α-GI:triazole (**4**) (crimson) with aspartic acids of the complementary δ **(A)** and γ **(B)** faces during nAChR binding. 1,5-Triazole bridge (gold), disulfide bond (yellow) and distance between interaction (dashed lines) (PDBID: 28WS).

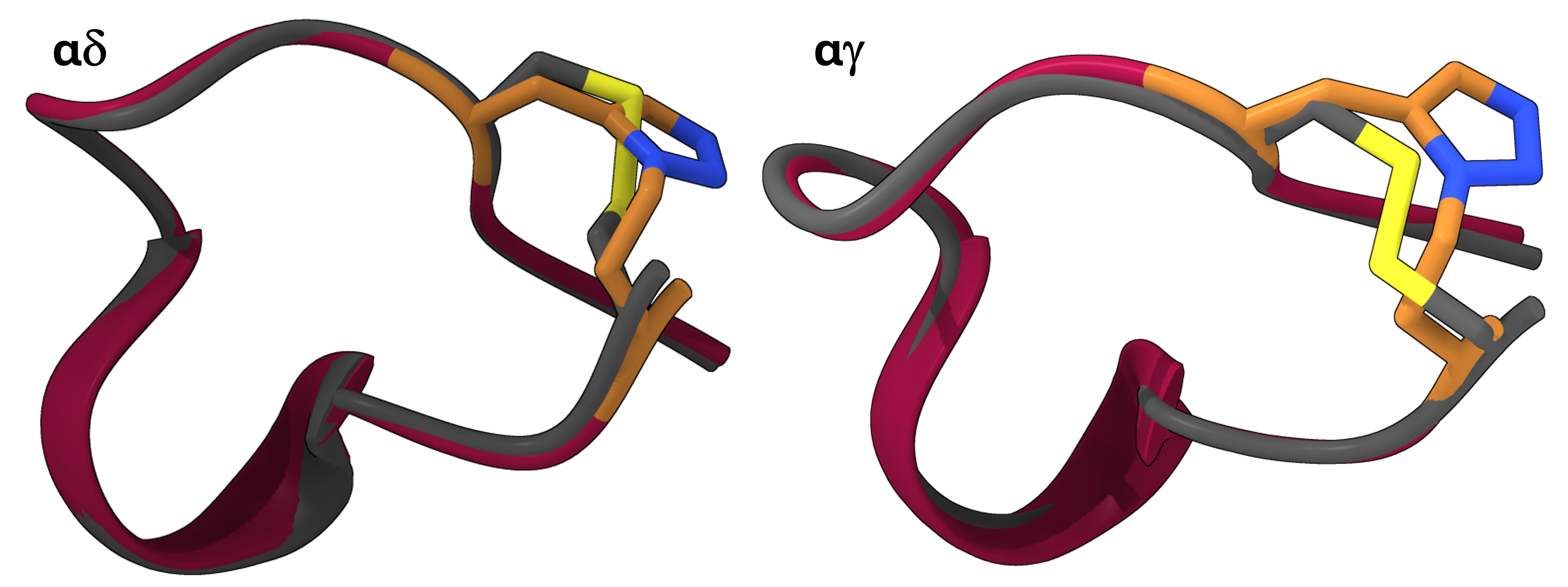

**Supporting Information Figure S11.** Overlay of bound disulfide α-GI peptide (**1**, grey) and 1,5-triazole α-GI peptidomimetic (**4**, crimson) at the two nAChR binding interfaces. Heteroatoms for the variable bridge residues coloured accordingly.

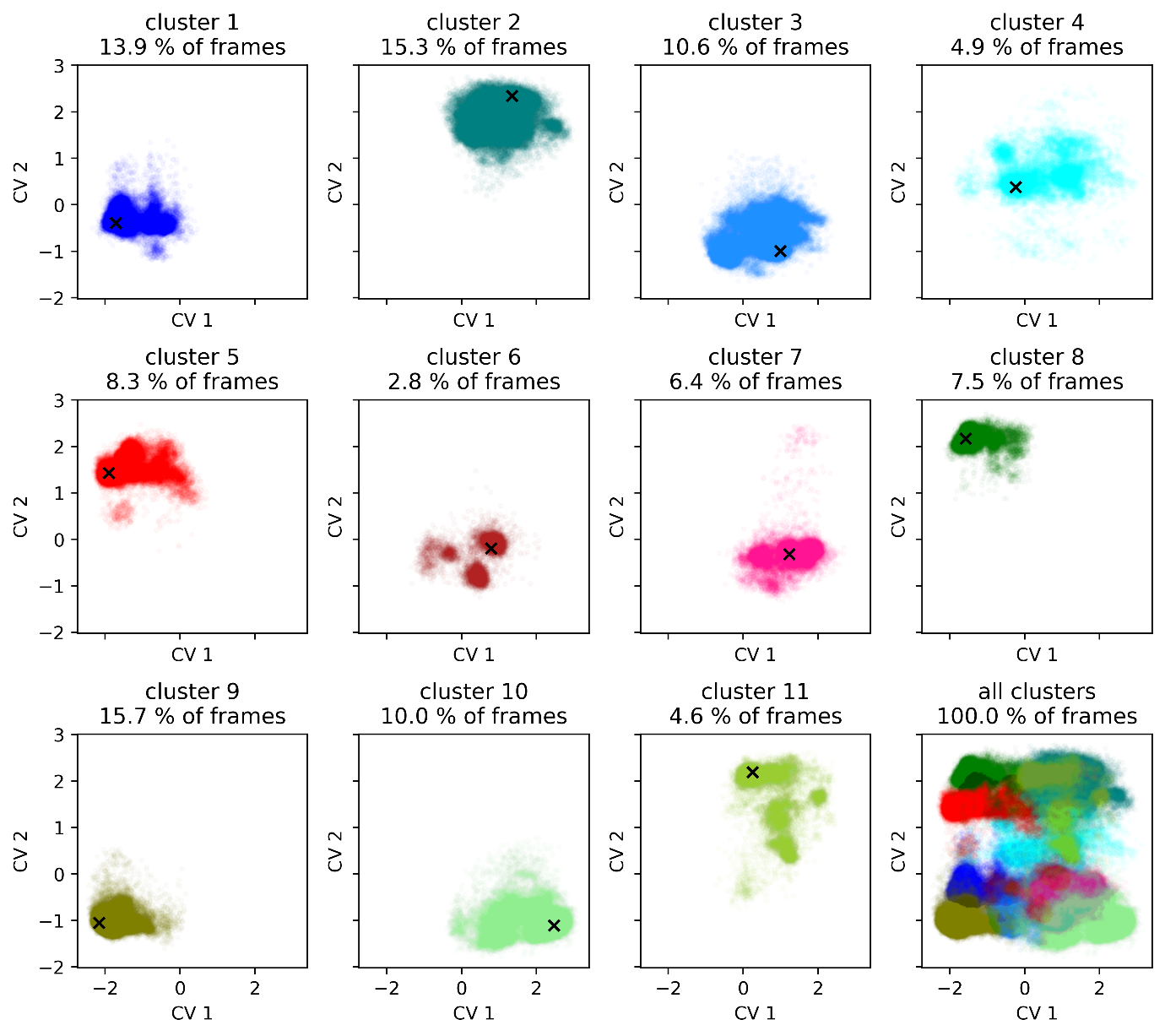

**Supporting Information Figure S12**. Result of k-means NANI clustering of the combined 1,4 and 1,5-triazole mimetics and wild type α-GI trajectories. Percentages of frames represent the cluster populations in relation to the combined trajectory and do not reflect the populations of each cluster in each separate trajectory (see **Table SX** for breakdown of cluster populations per trajectory). Each cluster has been projected back onto the principal component space of wild type α-GI as in Figure 1 and the bottom right plot shows the overlay of all 11 clusters reproducing the original PC plot. Cluster centroids are marked with a black cross and CV 1 and 2 relate back to PC1 and PC2 of the PCA plot. Cluster 4 (top right in light blue) neatly occupies the unique portion of the distribution originally noted.

**Supporting Information Table S1.** Cluster populations by compound.

| **Cluster number** | **1,4-triazole mimetic proportion (%)** | **1,5-triazole mimetic proportion (%)** | **Wild type α-GI proportion (%)** |
| --- | --- | --- | --- |
| **1** | 8.68 | 23.00 | 10.07 |
| **2** | 11.75 | 18.97 | 15.17 |
| **3** | 9.51 | 8.05 | 14.34 |
| **4** | 11.92 | 0.70 | 2.17 |
| **5** | 6.69 | 11.44 | 6.67 |
| **6** | 8.33 | 0.05 | 0.01 |
| **7** | 10.79 | 3.60 | 4.95 |
| **8** | 4.23 | 13.11 | 5.13 |
| **9** | 18.57 | 9.53 | 18.93 |
| **10** | 4.03 | 6.66 | 19.28 |
| **11** | 5.50 | 4.89 | 3.28 |

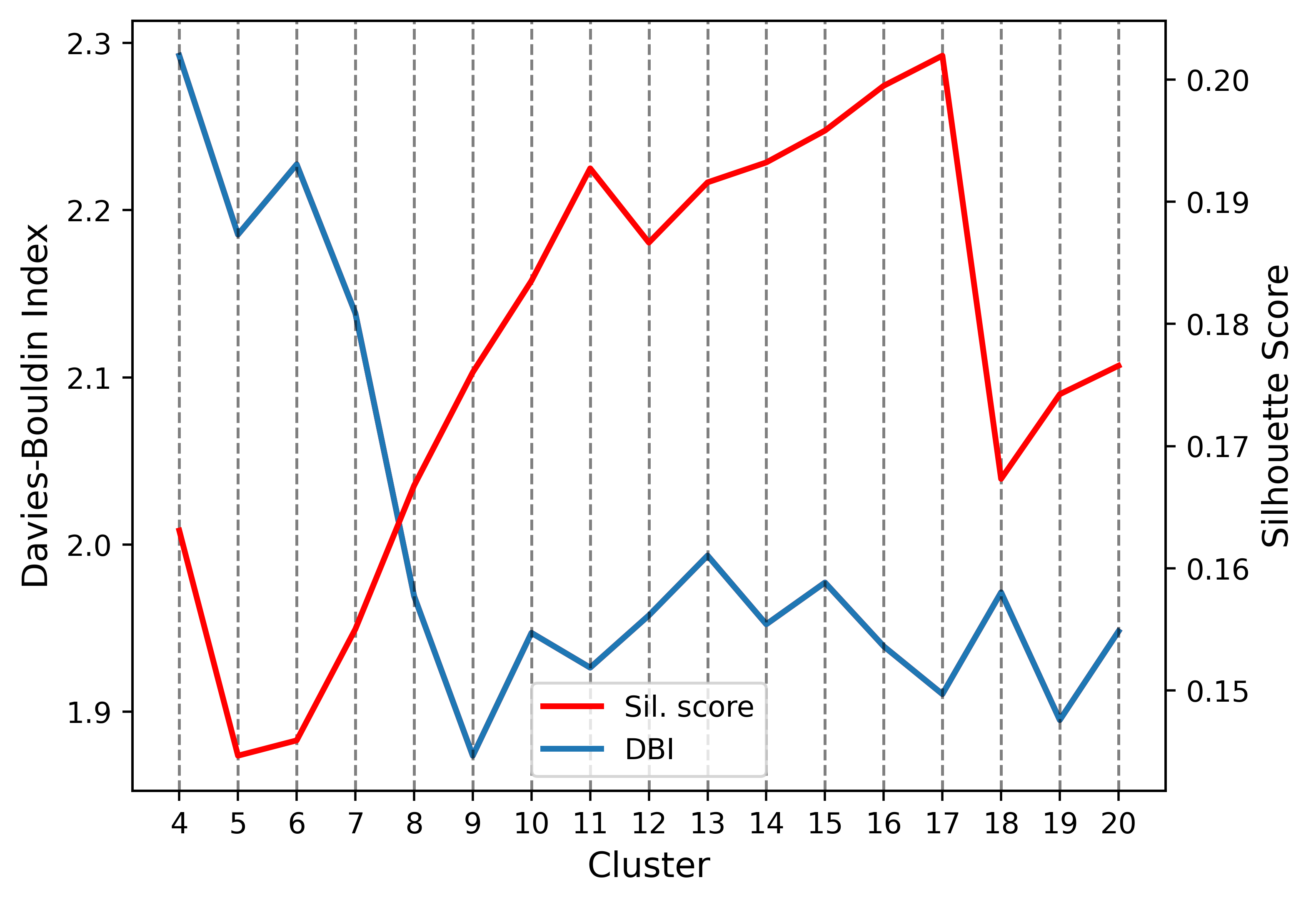

**Supporting Information Figure S13**. Elbow plot for the cluster centroid analysis of alpha-GI clustering on backbone dihedral angles. Maximizing silhouette score (red) while minimising DBI (blue) leads to cluster numbers of eleven or seventeen being optimal for this system. Eleven clusters were chosen as although seventeen showed slightly improved metrics, eleven is easier to handle for further analysis of the individual clusters.

### Experimental

#### Table of Peptides

**Supporting Information Table S2.** Name, sequence, % yield, % purity, *m/z* and retention time of peptides. Abbreviations: 3-BA = 3-butynoic acid, Dab = 2,3-diaminobutyric acid, Pra = propargylglycine, Aha = azidohomoalanine, Ac = acetyl, Ac-Cl = acetyl chloride, NH_2_ = C-terminal amide. Detailed characterisation data for peptides can be found in **Supporting Information Figures S14 – S31**.

| **Peptide** | **Sequence** | **Yield (%)** | **% Purity** | **Calculated *m/z*** | **Observed *m/z*** | **t_R_ (mins)** |
| --- | --- | --- | --- | --- | --- | --- |
| **1**^a^ | *H-ECCNPACGRHYTSC-NH_2_* | 7 | 99 | 1437.61 | 1437.80 ± 0.49 | 17.6 |
| **1**^a^ | *H-ECCNPACGRHYTSC-NH_2_* |  | 96 | 1437.67 | 1437.74 ± 0.23 | 17.1 |
| **2**^b^ | *H-EC(Pra)NPACGRHYTS(Aha)-NH_2_* | 2 | 96 | 1454.57 | 1454.13 ± 0.18 | 17.5 |
| **3**^b^ | *H-EC(Aha)NPACGRHYTS(Pra)-NH_2_* | 2 | 96 | 1454.57 | 1454.11 ± 0.21 | 16.8 |
| **4**^c^ | *H-EC(Pra)NPACGRHYTS(Aha)-NH_2_* | 2 | 99 | 1454.57 | 1454.09 ± 0.18 | 16.6 |
| **5**^c^ | *H-EC(Aha)NPACGRHYTS(Pra)-NH_2_* | 2 | 97 | 1454.57 | 1454.14 ± 0.11 | 17.6 |
| **6**^a^ | *H-ECCNPACGR(Dab[Ac])YTSC-NH_2_* | 2 | 98 | 1442.62 | 1442.51 ± 0.13 | 17.8 |
| **7**^a^ | *H-ECCNPACGR(Dab[3-BA])YTSC-NH_2_* | 2 | 95 | 1466.65 | 1466.02 ± 0.49 | 18.3 |
| **8**^a^ | *H-ECCNPACGR(Dab[Ac-Cl])YTSC-NH_2_* | 2 | 95 | 1477.07 | 1476.70 ± 0.43 | 18.1 |

^a^Peptide contains two disulfide bridges between cysteines 2/7 and 3/13 respectively.

^b^Peptide contains a disulfide bridge between cysteines 2/7 and a 1,4-triazole bridge between residues 3/13.

^c^Peptide contains a disulfide bridge between cysteines 2/7 and a 1,5-triazole bridges between residues 3/13.

#### Peptide Synthesis

##### Synthesis of Native α-GI (**1**) by Oxidative Folding

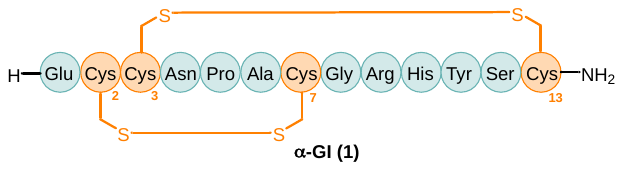

Linear α-GI (**1**) was synthesised by automated Fmoc-SPPS as described in **General Method 1A (0.1 mmol)** whereby all Cys residues were Trt protected. The linear peptide was liberated from the resin and its protecting groups simultaneously removed under the conditions described in **General Method 10A** to afford the crude linear peptide (100 mg, 70% yield [based on initial resin loading]) at approximately 65% purity. The crude linear peptide was cyclised as described in **General Method 11** to yield the desired cyclic peptide, **1**.

Crude peptide **1** (100 mg) was solubilised in 0.1% TFA in MeCN:MQ H_2_O (2:8, *v/v*) at a concentration of ~30 mg/mL and purified by preparative RP-HPLC (2 x 1700 μL injection) employing a gradient of 15% – 65%B over 50 min (*ca.* 1%B/min) at a flow rate of 10 mL/min. Fractions were analysed by RP-HPLC and ESI-MS for compound identification and lyophilised to afford the compound, α-GI (**1**), as a white amorphous powder (9.6 mg, 10% recovery [based on crude yield], 99% purity, 7% overall yield).

**ESI-MS**: Mass calculated for [C_55_H_80_N_20_O_18_S_4_ + H] 1437.61; deconvoluted mass observed: 1437.80 ± 0.49. Charge states; 480.45 [M+3H]^3+^, 719.82 [M+2H]^2+^, 1438.41 [M+H]^+^.

**RP-HPLC**: t_R_ = 17.6 min. Phenomenex Aeris Peptide XB-C18 (100 Å, 5 µm, 150 mm x 4.6 mm), linear gradient 5% – 95%B over 50 min (*ca.* 1.8%B/min) at 1 mL/min.

##### Synthesis of Native α-GI (**1**) by an Orthogonal Strategy

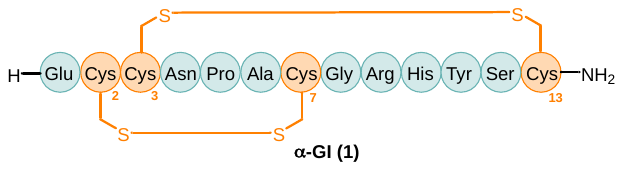

Linear α-GI (**1**) was synthesised by automated Fmoc-SPPS as described in **General Method 1B (0.1 mmol)** whereby Cys residues at positions 2 and 7 were Mmt protected and Cys residues at positions 3 and 13 were SIT protected. Disulfide bonds were formed asynchronously by **General Methods** **2** and **3**, followed by **General Methods 4 and 3** to form disulfide bonds in order between Cys^3/13^ followed by Cys^2/7^. The full folded peptide was liberated from the resin and its protecting groups simultaneously removed under the conditions described in **General Method 10B** to afford the crude peptide (9.08 mg, 6% yield [based on initial resin loading]) at approximately 70% purity.

Crude peptide **1** (9.08 mg) was solubilised in 0.1% TFA in MeCN:MQ H_2_O (2:8, *v/v*) at a concentration of ~9 mg/mL and purified by preparative RP-HPLC (1 x 1000 μL injection) employing a gradient of 15% – 65%B over 50 min (*ca.* 1%B/min) at a flow rate of 10 mL/min. Fractions were analysed by RP-HPLC and ESI-MS for compound identification and lyophilised to afford the compound, α-GI (**1**), as a white amorphous powder (1.0 mg, 11% recovery [based on crude yield], 96% purity, 1% overall yield).

**ESI-MS**: Mass calculated for [C_55_H_80_N_20_O_18_S_4_ + H] 1437.61; deconvoluted mass observed: 1437.74 ± 0.23. Charge states; 719.79 [M+2H]^2+^, 1438.90 [M+H] ^+^.

**RP-HPLC**: t_R_ = 17.1 min. Phenomenex Aeris Peptide XB-C18 (100 Å, 5 µm, 150 mm x 4.6 mm), linear gradient 5% – 95%B over 50 min (*ca.* 1.8%B/min) at 1 mL/min.

##### Synthesis of α-GI-1,4-triazole (**2**)

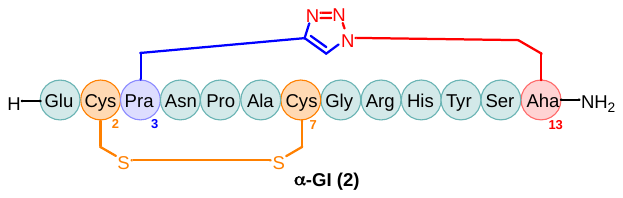

Linear peptide **2** was synthesised by automated Fmoc-SPPS as described in **General Method 1 (0.1 mmol)**. Following elongation of the linear peptide the resin was split into two portions of 0.05 mmol (to be split between peptides **2** and **4**). The linear peptide underwent a CuAAC as described in **General Method 6**. On-resin Acm removal and concomitant disulfide bond formation between Cys^2/7^ was mediated as described by **General Method 7**. The on-resin fully folded peptide was liberated from the by **General Method 10B** to afford the crude cyclic peptide (34 mg, 46% yield [based on initial resin loading]) at approximately 64% purity.

Crude peptide **2** (34 mg) was solubilised in 0.1% TFA in MeCN:MQ H_2_O (2:8, *v/v*) at a concentration of ~20 mg/mL and purified by preparative RP-HPLC (1 x 1700 μL injection) employing a gradient of 15% – 65%B over 50 min (*ca.* 1%B/min) at a flow rate of 10 mL/min. Fractions were analysed by RP-HPLC and ESI-MS for compound identification and lyophilised to afford the compound, **2**, as a white amorphous powder (1.2 mg, 4% recovery [based on crude yield], 96% purity, 2% overall yield).

**ESI-MS**: Mass calculated for [C_58_H_83_N_23_O_18_S_2_ + H] 1454.57; deconvoluted mass observed: 1454.13 ± 0.18. Charge states; 485.68 [M+3H]^3+^, 728.00 [M+2H]^2+^.

**RP-HPLC**: t_R_ = 17.5 min. Phenomenex Aeris Peptide XB-C18 (100 Å, 5 µm, 150 mm x 4.6 mm), linear gradient 5% – 95%B over 50 min (*ca.* 1.8%B/min) at 1 mL/min.

##### Synthesis of α-GI-1,4-triazole (**3**)

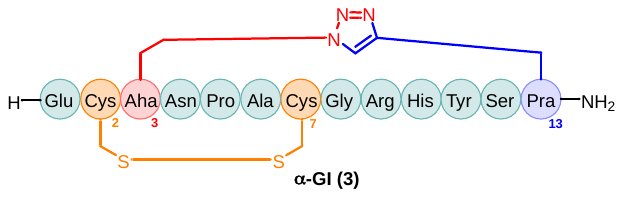

Linear peptide **3** was synthesised by automated Fmoc-SPPS as described in **General Method 1 (0.1 mmol)**. Following elongation of the linear peptide the resin was split into two portions of 0.05 mmol (to be split between peptides **3** and **5**). The linear peptide underwent a CuAAC as described in **General Method 6**. On-resin Acm removal and concomitant disulfide bond formation between Cys^2/7^ was mediated as described by **General Method 7**. The on-resin fully folded peptide was liberated from the by **General Method 10B** to afford the crude cyclic peptide (41 mg, 56% yield [based on initial resin loading]) at approximately 67% purity.

Crude peptide **3** (41 mg) was solubilised in 0.1% TFA in MeCN:MQ H_2_O (2:8, *v/v*) at a concentration of ~25 mg/mL and purified by preparative RP-HPLC (1 x 1700 μL injection) employing a gradient of 15% – 65%B over 50 min (*ca.* 1%B/min) at a flow rate of 10 mL/min. Fractions were analysed by RP-HPLC and ESI-MS for compound identification and lyophilised to afford the compound, **3**, as a white amorphous powder (1.6 mg, 4% recovery [based on crude yield], 96% purity, 2% overall yield).

**ESI-MS**: Mass calculated for [C_58_H_83_N_23_O_18_S_2_ + H] 1454.57; deconvoluted mass observed: 1454.11 ± 0.21. Charge states; 485.75 [M+3H]^3+^, 728.08 [M+2H]^2+^.

**RP-HPLC**: t_R_ = 16.8 min. Phenomenex Aeris Peptide XB-C18 (100 Å, 5 µm, 150 mm x 4.6 mm), linear gradient 5% – 95%B over 50 min (*ca.* 1.8%B/min) at 1 mL/min.

##### Synthesis of α-GI-1,5-triazole (**4**)

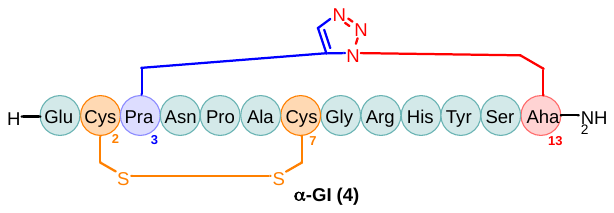

Linear peptide **4** was synthesised by automated Fmoc-SPPS as described in **General Method 1 (0.1 mmol)**. Following elongation of the linear peptide the resin was split into two portions of 0.05 mmol (to be split between peptides **2** and **4**). The linear peptide underwent a RuAAC as described in **General Method 8**. On-resin Acm removal and concomitant disulfide bond formation between Cys^2/7^ was mediated as described by **General Method 7**. The on-resin fully folded peptide was liberated from the by **General Method 10B** to afford the crude cyclic peptide (21 mg, 28% yield [based on initial resin loading]) at approximately 80% purity.

Crude peptide **4** (21 mg) was solubilised in 0.1% TFA in MeCN:MQ H_2_O (2:8, *v/v*) at a concentration of ~20 mg/mL and purified by preparative RP-HPLC (1 x 1100 μL injection) employing a gradient of 15% – 65%B over 50 min (*ca.* 1%B/min) at a flow rate of 10 mL/min. Fractions were analysed by RP-HPLC and ESI-MS for compound identification and lyophilised to afford the compound, **4**, as a white amorphous powder (1.2 mg, 6% recovery [based on crude yield], 99% purity, 2% overall yield).

**ESI-MS**: Mass calculated for [C_58_H_83_N_23_O_18_S_2_ + H] 1454.57; deconvoluted mass observed: 1454.09 ± 0.18. Charge states; 485.69 [M+3H]^3+^, 727.98 [M+2H]^2+^.

**RP-HPLC**: t_R_ = 16.6 min. Phenomenex Aeris Peptide XB-C18 (100 Å, 5 µm, 150 mm x 4.6 mm), linear gradient 5% – 95%B over 50 min (*ca.* 1.8%B/min) at 1 mL/min.

##### Synthesis of α-GI-1,5-triazole (**5**)

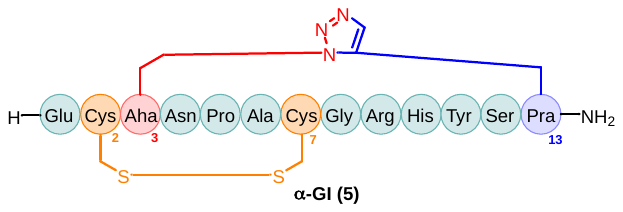

Linear peptide **5** was synthesised by automated Fmoc-SPPS as described in **General Method 1 (0.1 mmol)**. Following elongation of the linear peptide the resin was split into two portions of 0.05 mmol (to be split between peptides **3** and **5**). The linear peptide underwent a RuAAC as described in **General Method 8**. On-resin Acm removal and concomitant disulfide bond formation between Cys^2/7^ was mediated as described by **General Method 7**. The on-resin fully folded peptide was liberated from the by **General Method 10B** to afford the crude cyclic peptide (22 mg, 30% yield [based on initial resin loading]) at approximately 49% purity.

Crude peptide **5** (22 mg) was solubilised in 0.1% TFA in MeCN:MQ H_2_O (2:8, *v/v*) at a concentration of ~20 mg/mL and purified by preparative RP-HPLC (1 x 1000 μL injection) employing a gradient of 15% – 65%B over 50 min (*ca.* 1%B/min) at a flow rate of 10 mL/min. Fractions were analysed by RP-HPLC and ESI-MS for compound identification and lyophilised to afford the compound, **5**, as a white amorphous powder (2.1 mg, 9% recovery [based on crude yield], 97% purity, 2% overall yield).

**ESI-MS**: Mass calculated for [C_58_H_83_N_23_O_18_S_2_ + H] 1454.57; deconvoluted mass observed: 1454.14 ± 0.11. Charge states; 485.74 [M+3H]^3+^, 728.03 [M+2H]^2+^.

**RP-HPLC**: t_R_ = 17.6 min. Phenomenex Aeris Peptide XB-C18 (100 Å, 5 µm, 150 mm x 4.6 mm), linear gradient 5% – 95%B over 50 min (*ca.* 1.8%B/min) at 1 mL/min.

##### Synthesis of α-GI – H10Dab-Ac (**6**)

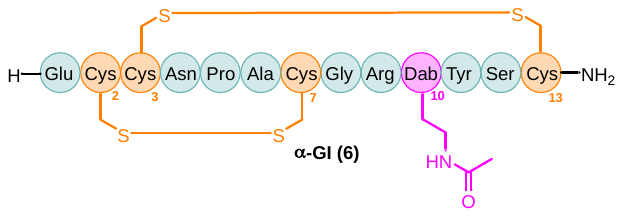

Linear peptide **6** was synthesised by automated Fmoc-SPPS as described in **General Method 1B (0.1 mmol)** whereby Cys residues at positions 2 and 7 were Mmt protected and Cys residues at positions 3 and 13 were SIT protected. Following complete elongation of the peptide the N-terminal Fmoc group was exchange for Boc as described by **General Method 5**. Disulfide bonds were formed asynchronously by **General Methods** **2** and **3**, followed by **General Methods 4 and 3** to form disulfide bonds in order between Cys3/13 followed by Cys2/7. Orthogonally protected Dab underwent Dde removal as described in **General Method 9**. The *N^γ^*-unprotected Dab residue underwent capping with a solution of Ac_2_O (47.2 µL, 0.5 mmol, 5 equiv.) and DIPEA (174.8 µL, 1.0 mmol, 10 equiv.) in DMF (10 mL) for 2 h, following which the resin was thorough washed with DMF, followed by CH_2_Cl_2_, and dried under vacuum. The fully functionalised peptide was liberated from the resin and its protecting groups simultaneously removed under the conditions described in **General Method 10B** to afford the crude peptide (28 mg, 40% yield [based on initial resin loading]) at approximately 48% purity.

Crude peptide **6** (28 mg) was solubilised in 0.1% TFA in MeCN:MQ H_2_O (2:8, *v/v*) at a concentration of ~20 mg/mL and purified by preparative RP-HPLC (1 x 1400 μL injection) employing a gradient of 15% – 65%B over 50 min (*ca.* 1%B/min) at a flow rate of 10 mL/min. Fractions were analysed by RP-HPLC and ESI-MS for compound identification and lyophilised to afford the compound, **6**, as a white amorphous powder (2.0 mg, 7% recovery [based on crude yield], 98% purity, 2% overall yield).

**ESI-MS**: Mass calculated for [C_55_H_83_N_19_O_19_S_4_ + H] 1442.62; deconvoluted mass observed: 1442.51 ± 0.13. Charge states; 722.30 [M+2H]^2+^, 1443.42 [M+H] ^+^.

**RP-HPLC**: t_R_ = 17.8 min. Phenomenex Aeris Peptide XB-C18 (100 Å, 5 µm, 150 mm x 4.6 mm), linear gradient 5% – 95%B over 50 min (*ca.* 1.8%B/min) at 1 mL/min.

##### Synthesis of α-GI – H10Dab-3BA (**7**)

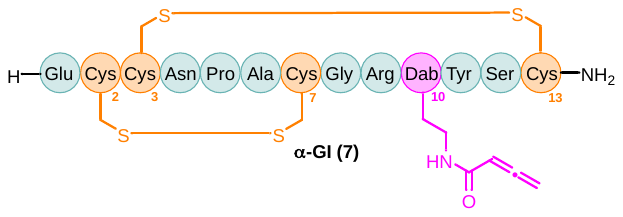

Linear peptide **7** was synthesised by automated Fmoc-SPPS as described in **General Method 1B (0.1 mmol)** whereby Cys residues at positions 2 and 7 were Mmt protected and Cys residues at positions 3 and 13 were SIT protected. Disulfide bonds were formed asynchronously by **General Methods** **2** and **3**, followed by **General Methods 4 and 3** to form disulfide bonds in order between Cys^3/13^ followed by Cys^2/7^. Orthogonally protected Dab underwent Dde removal as described in **General Method 9**. The *N^γ^*-unprotected Dab residue underwent capping with a solution of 3-BA (84.1 mg, 1.0 mmol, 10 equiv.), EEDQ (234.9 mg, 0.95 mmol, 9.5 equiv.) and *sym*-collidine (118.9 µL, 0.9 mmol, 9 equiv.) in CH_2_Cl_2_ (10 mL) for 18 h, following which the resin was thoroughly washed with DMF, followed by CH_2_Cl_2_, and dried under vacuum. The fully functionalised peptide was liberated from the resin and its protecting groups simultaneously removed under the conditions described in **General Method 10B** to afford the crude peptide (35 mg, 48% yield [based on initial resin loading]) at approximately 28% purity.

Crude peptide **7**(35 mg) was solubilised in 0.1% TFA in MeCN:MQ H_2_O (2:8, *v/v*) at a concentration of ~20 mg/mL and purified by preparative RP-HPLC (1 x 1700 μL injection) employing a gradient of 15% – 65%B over 50 min (*ca.* 1%B/min) at a flow rate of 10 mL/min. Fractions were analysed by RP-HPLC and ESI-MS for compound identification and lyophilised to afford the compound, **7**, as a white amorphous powder (3.0 mg, 9% recovery [based on crude yield], 95% purity, 4% overall yield).

**ESI-MS**: Mass calculated for [C_57_H_83_N_19_O_19_S_4_ + H] 1466.65; deconvoluted mass observed: 1466.02 ± 0.49. Charge states; 734.18 [M+2H]^2+^, 1466.67 [M+H] ^+^.

**RP-HPLC**: t_R_ = 18.3 min. Phenomenex Aeris Peptide XB-C18 (100 Å, 5 µm, 150 mm x 4.6 mm), linear gradient 5% – 95%B over 50 min (*ca.* 1.8%B/min) at 1 mL/min.

##### Synthesis of α-GI – H10Dab-Ac-Cl (**8**)

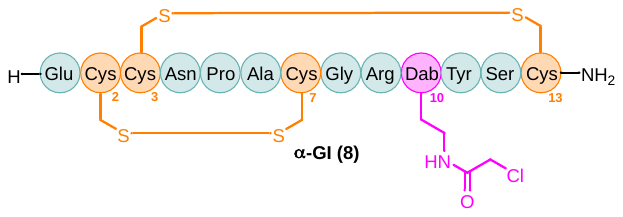

Linear peptide **8** was synthesised by automated Fmoc-SPPS as described in **General Method 1B (0.1 mmol)** whereby Cys residues at positions 2 and 7 were Mmt protected and Cys residues at positions 3 and 13 were SIT protected. Disulfide bonds were formed asynchronously by **General Methods** **2** and **3**, followed by **General Methods 4 and 3** to form disulfide bonds in order between Cys^3/13^ followed by Cys^2/7^. Orthogonally protected Dab underwent Dde removal as described in **General Method 9**. The *N^γ^*-unprotected Dab residue underwent capping with a solution of chloroacetyl chloride (79.5 µL, 1.0 mmol, 10 equiv.) and DIPEA (174.8 µL, 1.0 mmol, 10 equiv.) in DMF (10 mL) for 2 h, following which the resin was thorough washed with DMF, followed by CH_2_Cl_2_, and dried under vacuum. The fully functionalised peptide was liberated from the resin and its protecting groups simultaneously removed under the conditions described in **General Method 10B** to afford the crude peptide (29 mg, 40% yield [based on initial resin loading]) at approximately 45% purity.

Crude peptide **8** (29 mg) was solubilised in 0.1% TFA in MeCN:MQ H_2_O (2:8, *v/v*) at a concentration of ~20 mg/mL and purified by preparative RP-HPLC (1 x 1500 μL injection) employing a gradient of 15% – 65%B over 50 min (*ca.* 1%B/min) at a flow rate of 10 mL/min. Fractions were analysed by RP-HPLC and ESI-MS for compound identification and lyophilised to afford the compound, **8**, as a white amorphous powder (1.1 mg, 4% recovery [based on crude yield], 95% purity, 2% overall yield).

**ESI-MS**: Mass calculated for [C_55_H_82_ClN_19_O_19_S_4_ + H] 1477.07; deconvoluted mass observed: 1476.70 ± 0.43. Charge states; 739.50 [M+2H]^2+^, 1477.39 [M+H] ^+^.

**RP-HPLC**: t_R_ = 18.1 min. Phenomenex Aeris Peptide XB-C18 (100 Å, 5 µm, 150 mm x 4.6 mm), linear gradient 5% – 95%B over 50 min (*ca.* 1.8%B/min) at 1 mL/min.

##### Characterisation

###### Reverse Phase – High Performance Liquid Chromatography

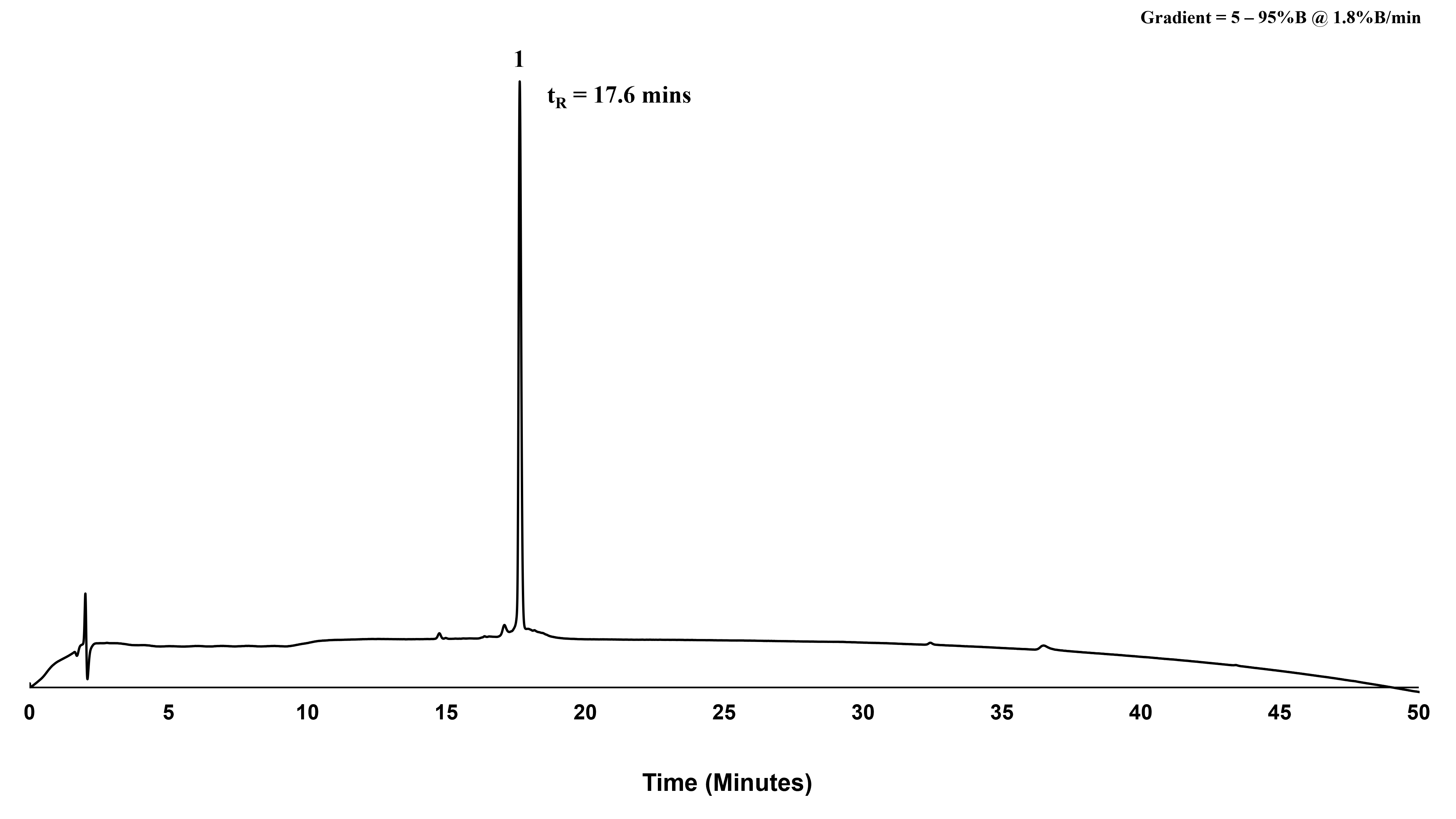

**Supporting Information Figure S14.** Analytical RP-HPLC chromatogram (214 nm) of purified peptide, **1 (by oxidative folding)** (*ca.* 99%) as analysed by peak area. Phenomenex Aeris Peptide XB-C18 (100 Å, 5 µm, 150 mm x 4.6 mm), linear gradient 5% – 95%B over 50 min (*ca.* 1.8%B/min) at 1 mL/min. t_R_ = 17.6 mins.

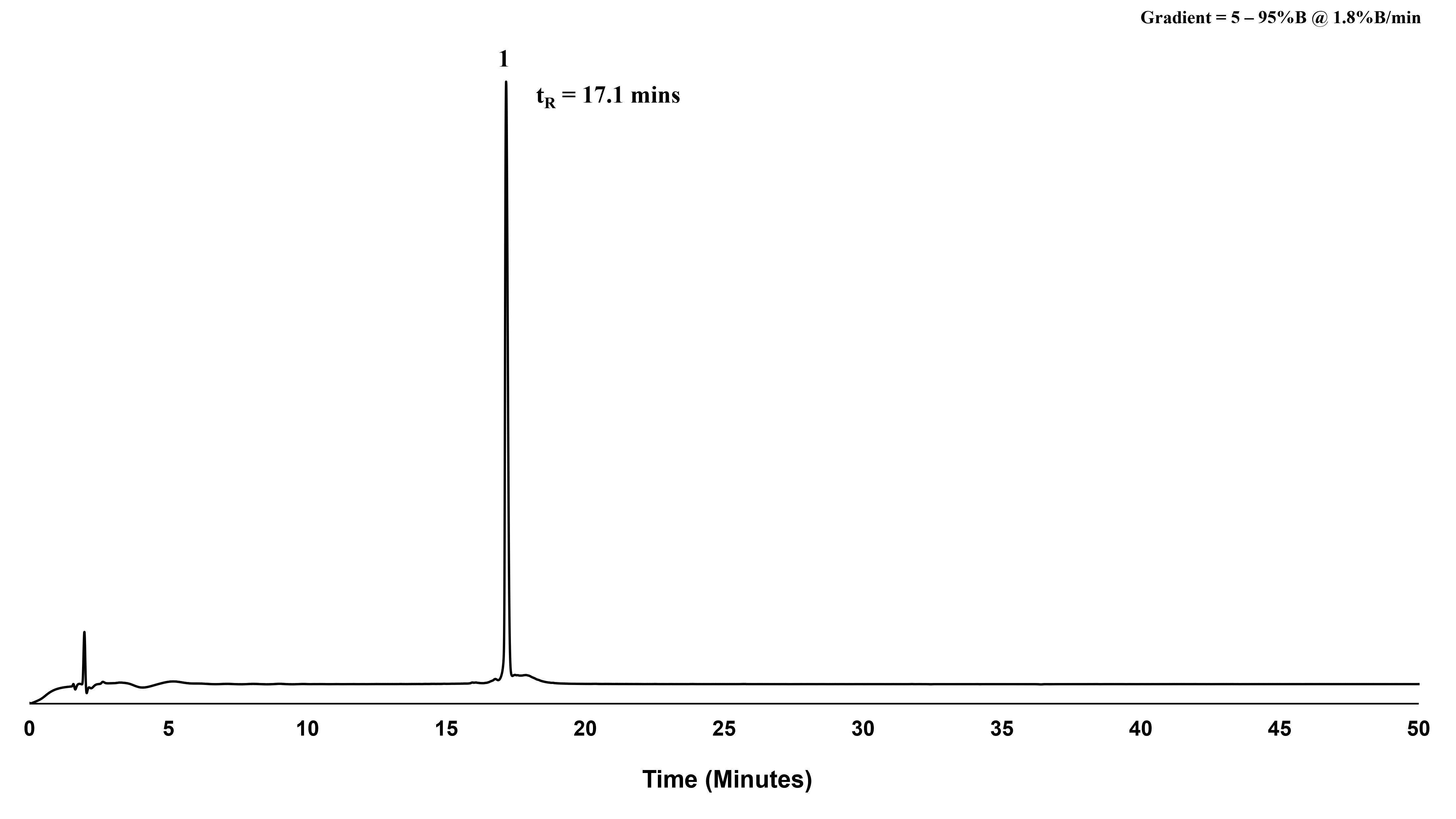

**Supporting Information Figure S15.** Analytical RP-HPLC chromatogram (214 nm) of purified peptide, **1 (by orthogonal synthesis)** (*ca.* 96%) as analysed by peak area. Phenomenex Aeris Peptide XB-C18 (100 Å, 5 µm, 150 mm x 4.6 mm), linear gradient 5% – 95%B over 50 min (*ca.* 1.8%B/min) at 1 mL/min. t_R_ = 17.1 mins.

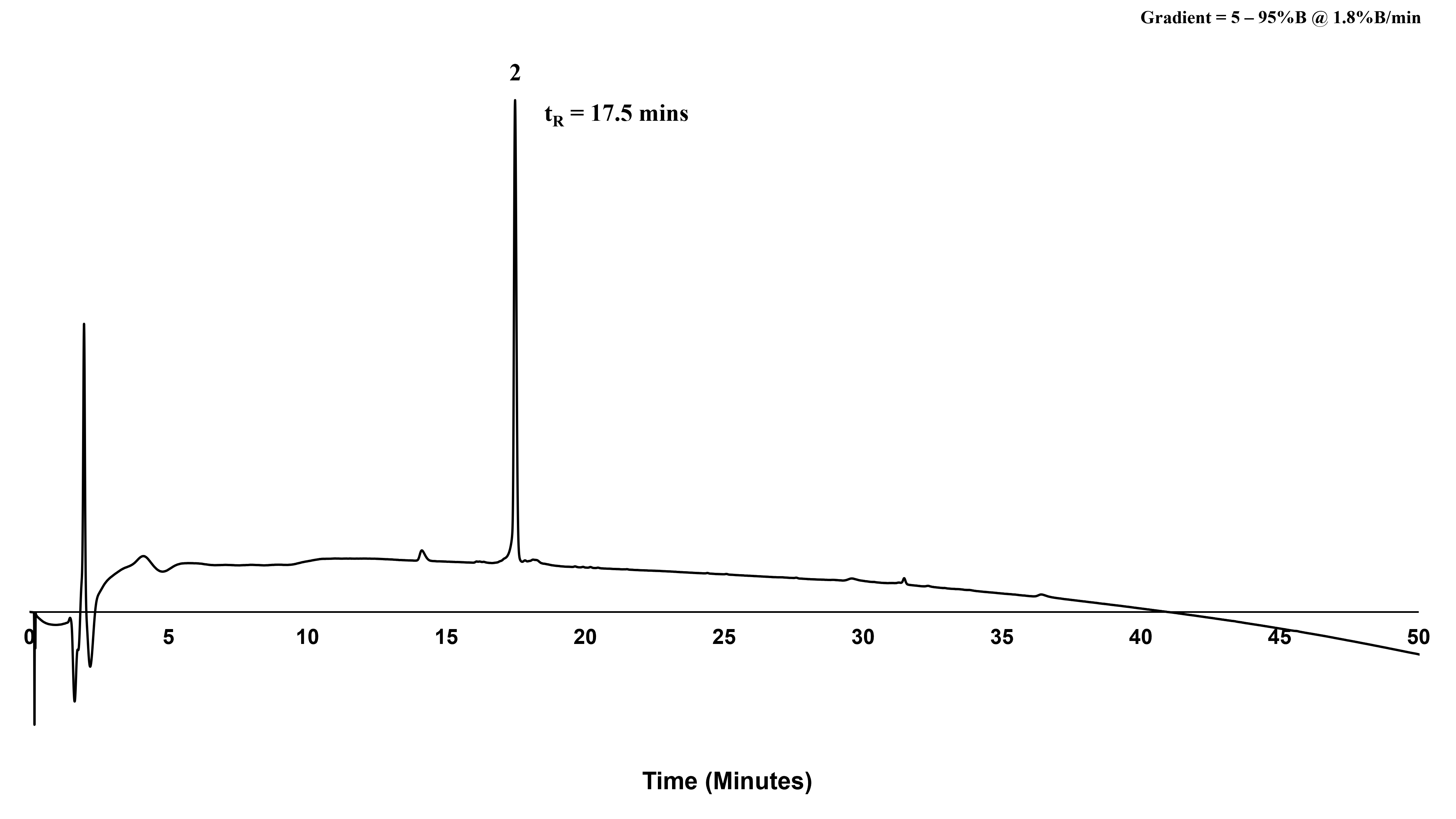

**Supporting Information Figure S16.** Analytical RP-HPLC chromatogram (214 nm) of purified peptide, **2** (*ca.* 96%) as analysed by peak area. Phenomenex Aeris Peptide XB-C18 (100 Å, 5 µm, 150 mm x 4.6 mm), linear gradient 5% – 95%B over 50 min (*ca.* 1.8%B/min) at 1 mL/min. t_R_ = 17.5 mins.

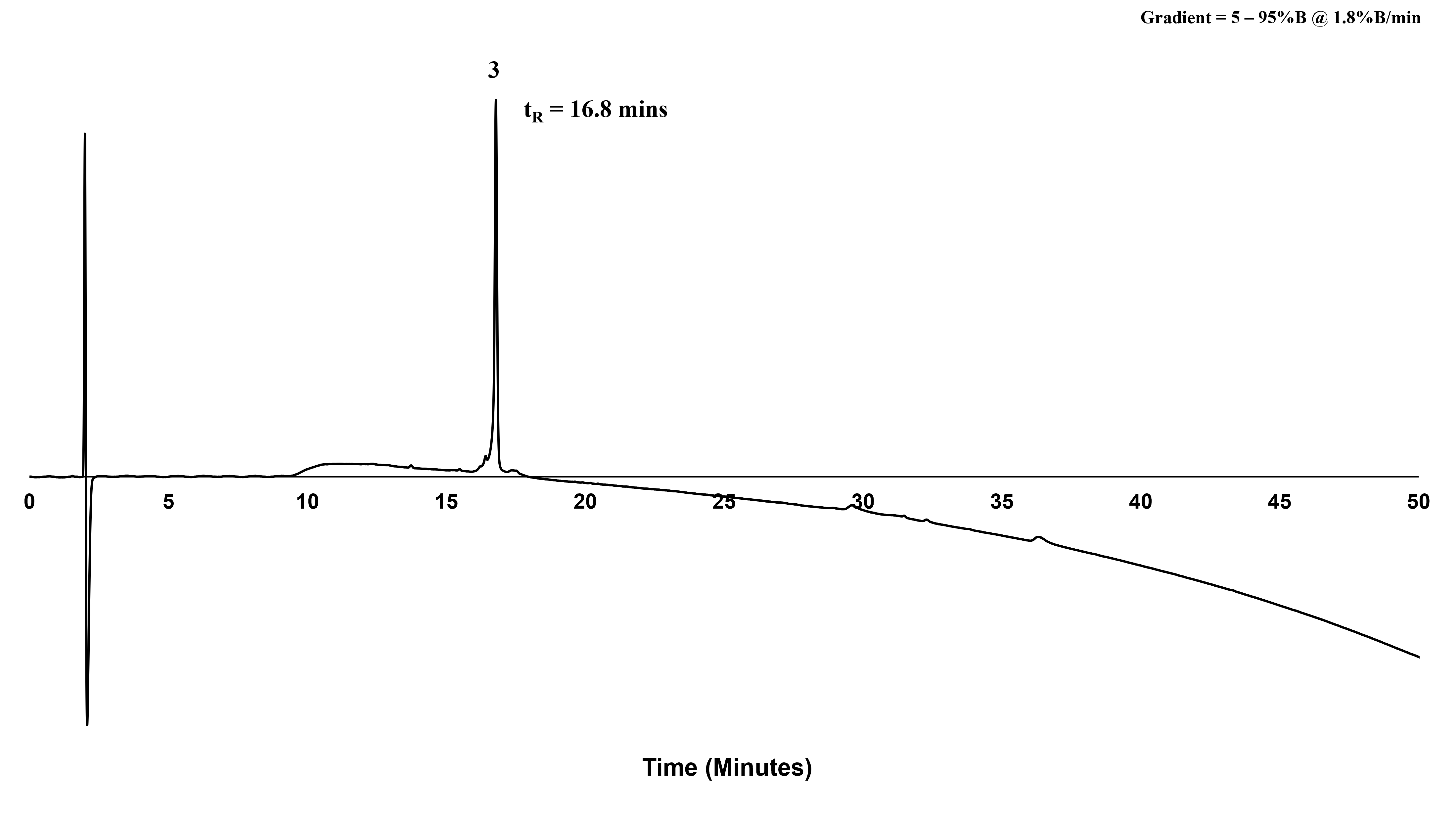

**Supporting Information Figure S17.** Analytical RP-HPLC chromatogram (214 nm) of purified peptide, **3** (*ca.* 96%) as analysed by peak area. Phenomenex Aeris Peptide XB-C18 (100 Å, 5 µm, 150 mm x 4.6 mm), linear gradient 5% – 95%B over 50 min (*ca.* 1.8%B/min) at 1 mL/min. t_R_ = 16.8 mins.

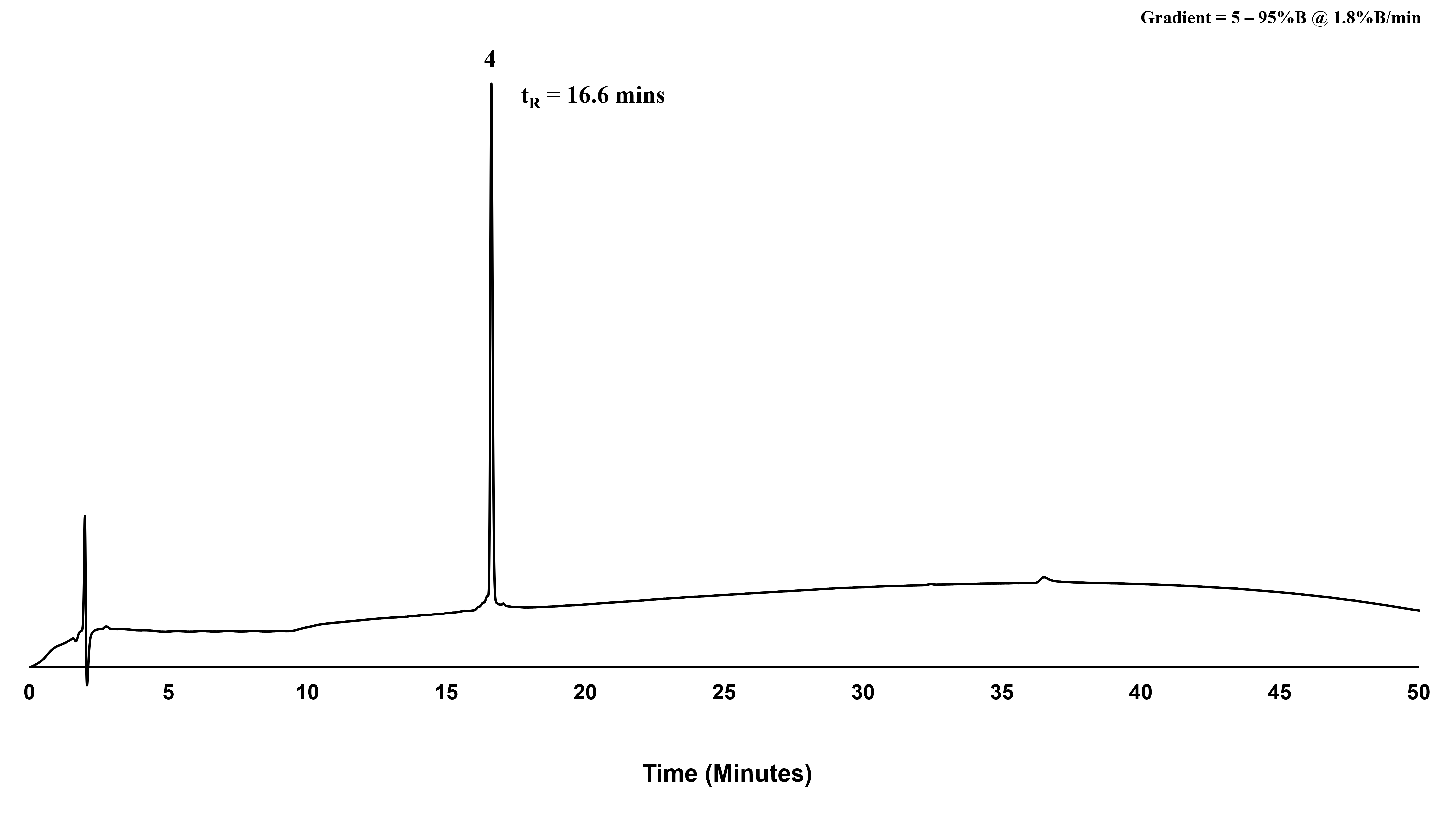

**Supporting Information Figure S18.** Analytical RP-HPLC chromatogram (214 nm) of purified peptide, **4** (*ca.* 99%) as analysed by peak area. Phenomenex Aeris Peptide XB-C18 (100 Å, 5 µm, 150 mm x 4.6 mm), linear gradient 5% – 95%B over 50 min (*ca.* 1.8%B/min) at 1 mL/min. t_R_ = 16.6 mins.

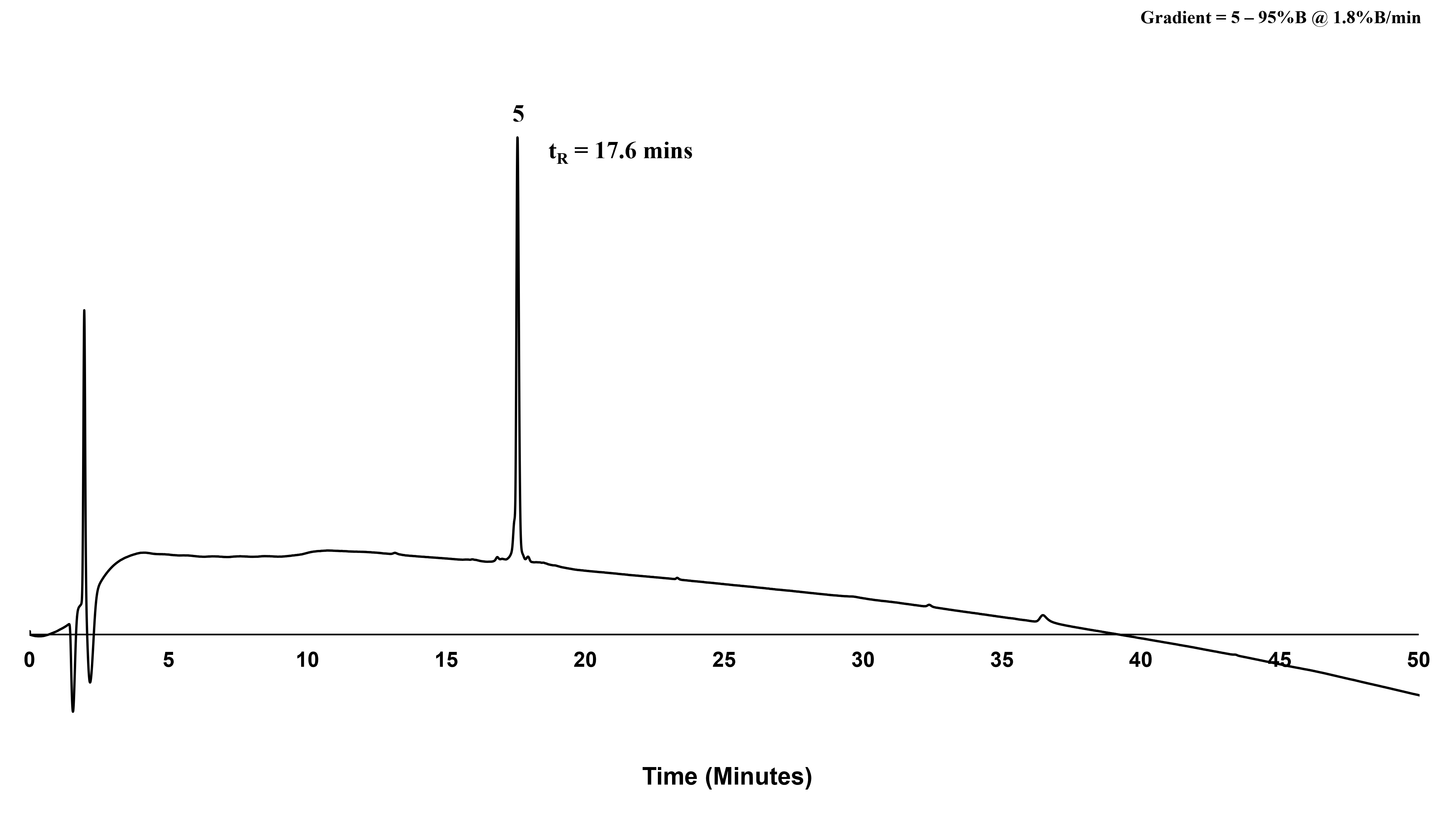

**Supporting Information Figure S19.** Analytical RP-HPLC chromatogram (214 nm) of purified peptide, **5** (*ca.* 97%) as analysed by peak area. Phenomenex Aeris Peptide XB-C18 (100 Å, 5 µm, 150 mm x 4.6 mm), linear gradient 5% – 95%B over 50 min (*ca.* 1.8%B/min) at 1 mL/min. t_R_ = 17.6 mins.

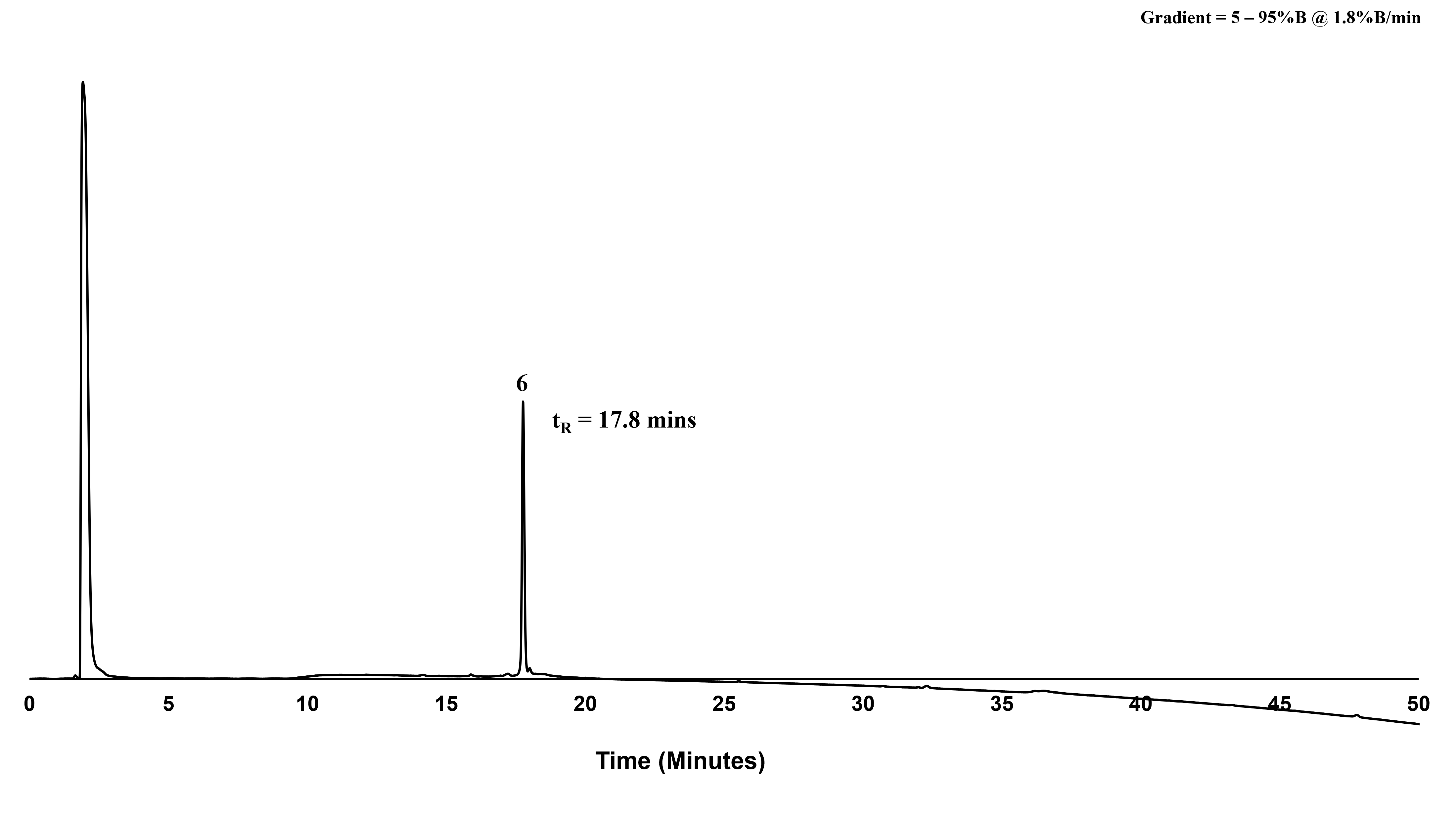

**Supporting Information Figure S20.** Analytical RP-HPLC chromatogram (214 nm) of purified peptide, **6** (*ca.* 98%) as analysed by peak area. Phenomenex Aeris Peptide XB-C18 (100 Å, 5 µm, 150 mm x 4.6 mm), linear gradient 5% – 95%B over 50 min (*ca.* 1.8%B/min) at 1 mL/min. t_R_ = 17.8 mins.

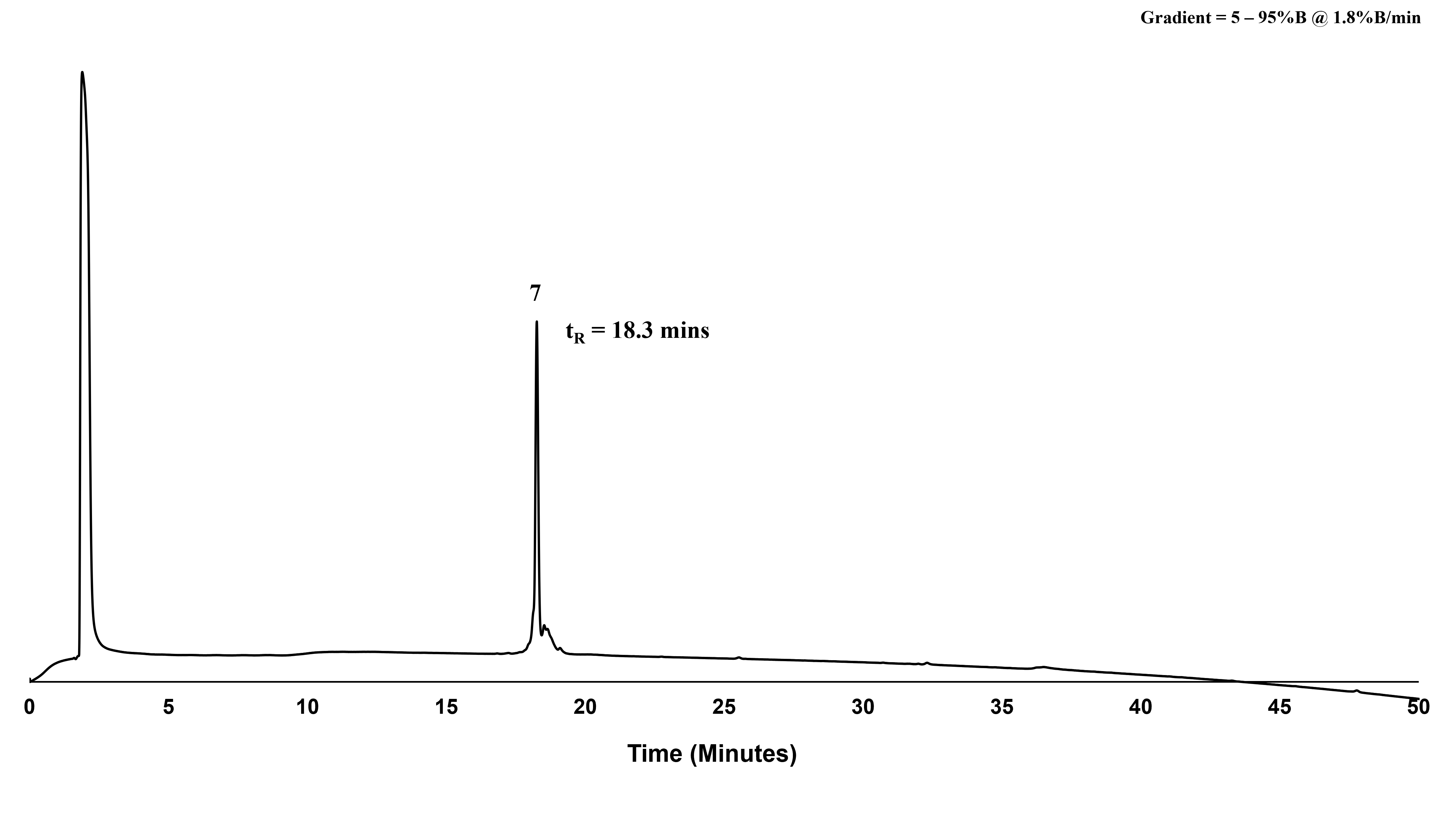

**Supporting Information Figure S21.** Analytical RP-HPLC chromatogram (214 nm) of purified peptide, **7** (*ca.* 95%) as analysed by peak area. Phenomenex Aeris Peptide XB-C18 (100 Å, 5 µm, 150 mm x 4.6 mm), linear gradient 5% – 95%B over 50 min (*ca.* 1.8%B/min) at 1 mL/min. t_R_ = 18.3 mins.

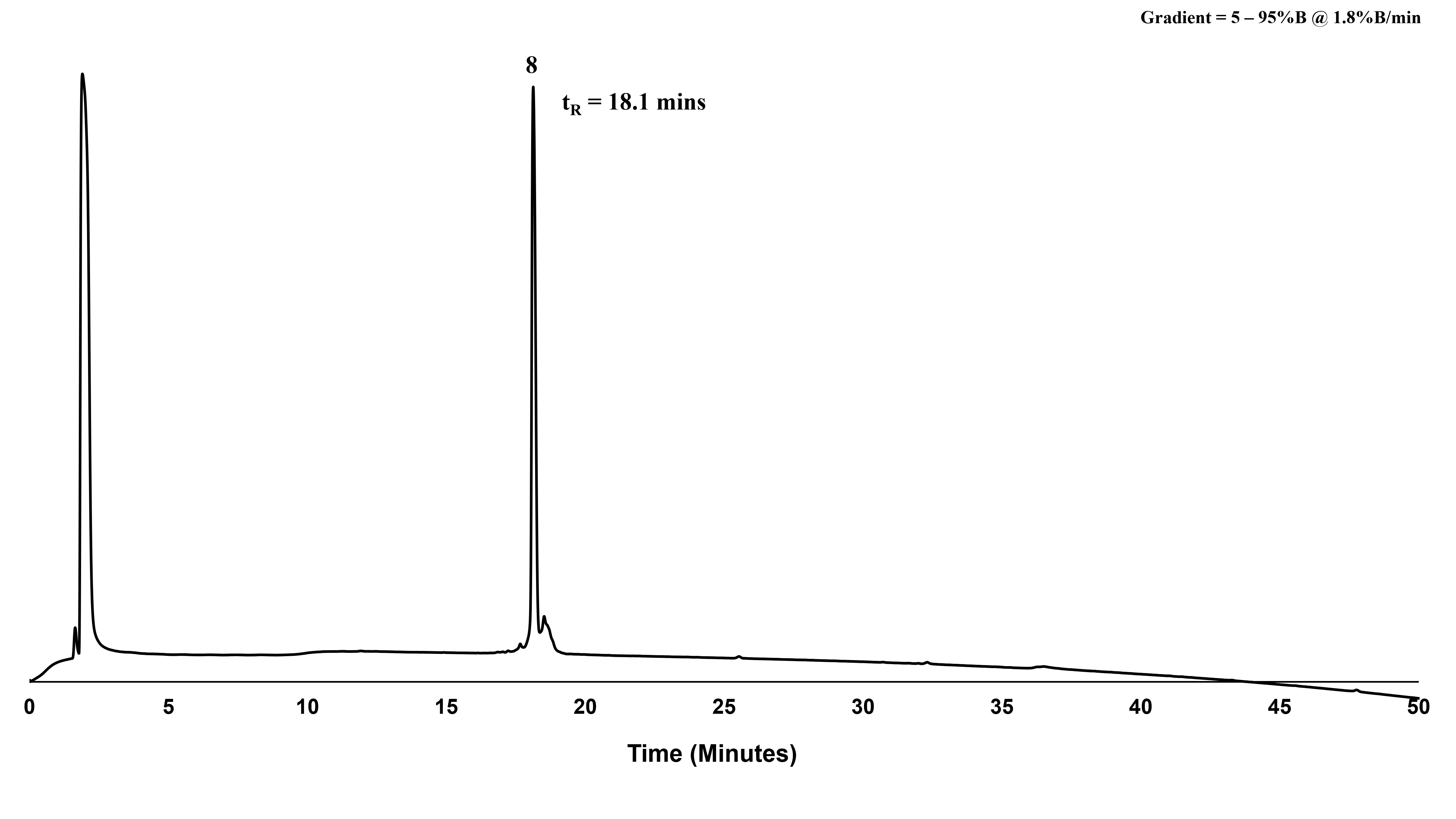

**Supporting Information Figure S22.** Analytical RP-HPLC chromatogram (214 nm) of purified peptide, **8** (*ca.* 95%) as analysed by peak area. Phenomenex Aeris Peptide XB-C18 (100 Å, 5 µm, 150 mm x 4.6 mm), linear gradient 5% – 95%B over 50 min (*ca.* 1.8%B/min) at 1 mL/min. t_R_ = 18.1 mins.

###### Liquid Chromatography Mass Spectrometry (ESI-MS)

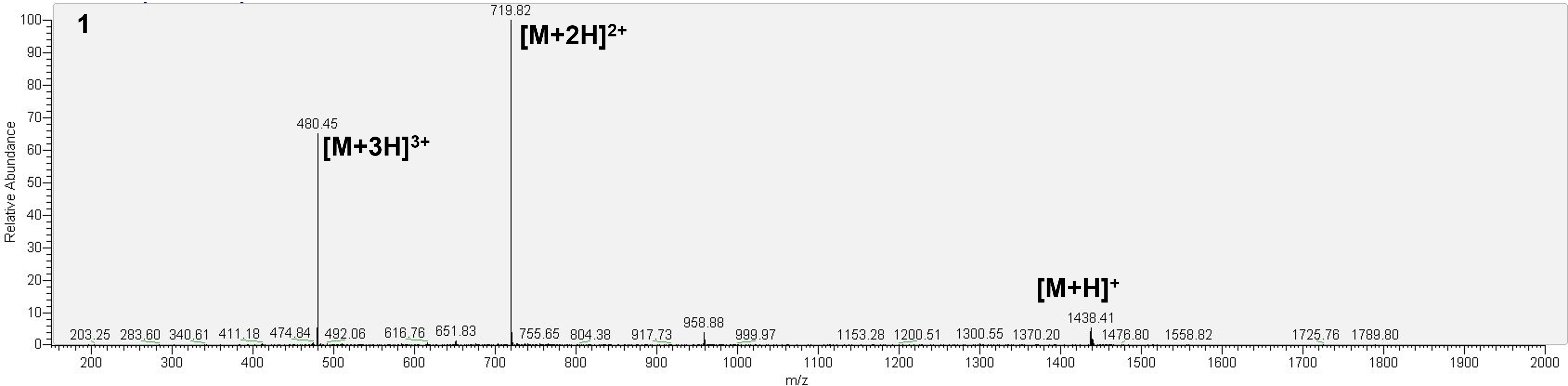

**Supporting Information Figure S23.** ESI-MS of purified peptide, **1 (by oxidative folding)**, mass calculated for [C_55_H_80_N_20_O_18_S_4_ + H] 1437.61; deconvoluted mass observed: 1437.80 ± 0.49. Charge states; 480.45 [M+3H]^3+^, 719.82 [M+2H]^2+^, 1438.41 [M+H]^+^.

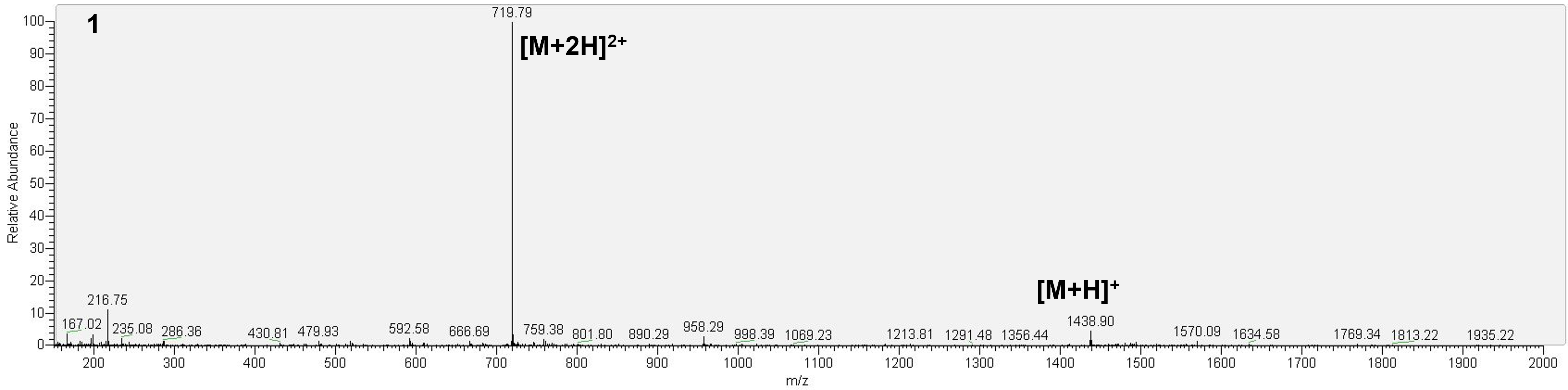

**Supporting Information Figure S24.** ESI-MS of purified peptide, **1 (by orthogonal synthesis)**, mass calculated for [C_55_H_80_N_20_O_18_S_4_ + H] 1437.61; deconvoluted mass observed: 1437.74 ± 0.23. Charge states; 719.79 [M+2H]^2+^, 1438.90 [M+H] ^+^.

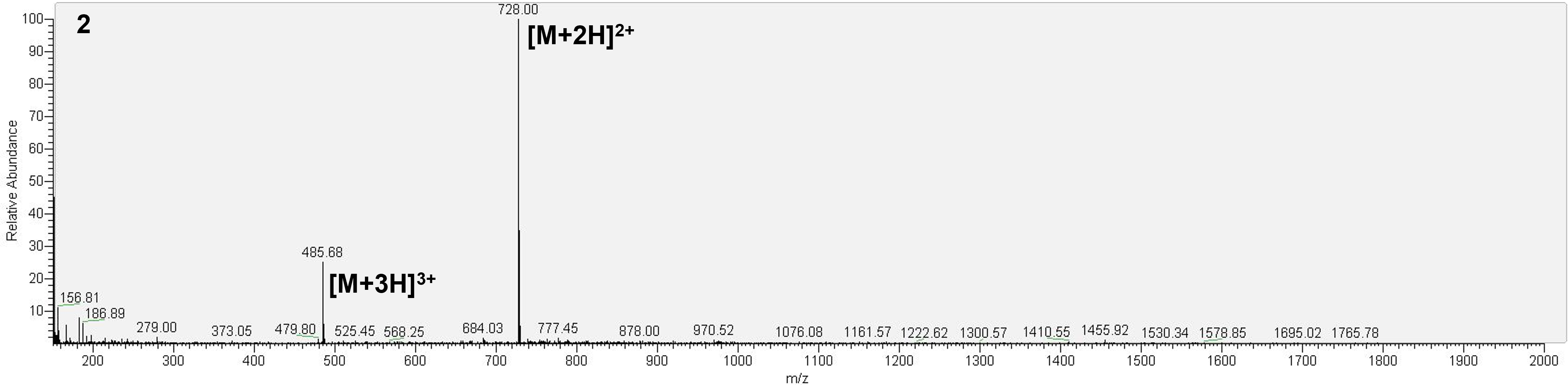

**Supporting Information Figure S25.** ESI-MS of purified peptide, **2**, mass calculated for [C_58_H_83_N_23_O_18_S_2_ + H] 1454.57; deconvoluted mass observed: 1454.13 ± 0.18. Charge states; 485.68 [M+3H]^3+^, 728.00 [M+2H]^2+^.

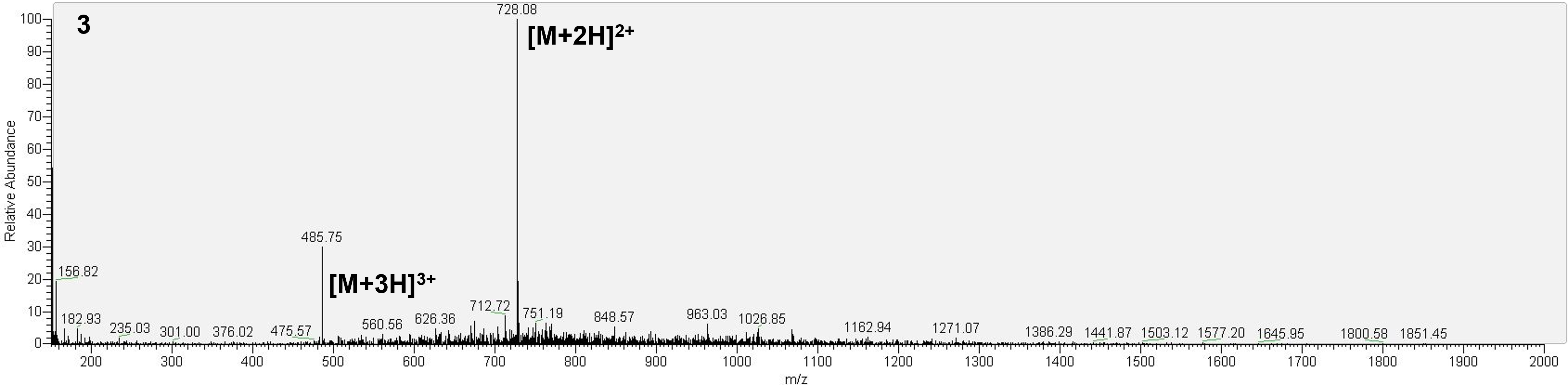

**Supporting Information Figure S26.** ESI-MS of purified peptide, **3**, mass calculated for [C_58_H_83_N_23_O_18_S_2_ + H] 1454.57; deconvoluted mass observed: 1454.11 ± 0.21. Charge states; 485.75 [M+3H]^3+^, 728.08 [M+2H]^2+^.

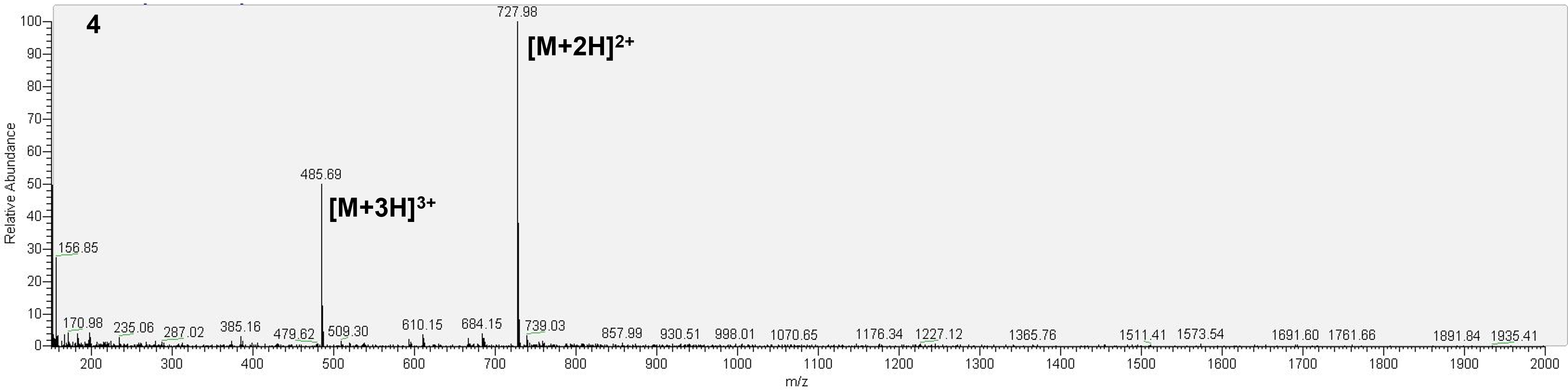

**Supporting Information Figure S27.** ESI-MS of purified peptide, **4**, mass calculated for [C_58_H_83_N_23_O_18_S_2_ + H] 1454.57; deconvoluted mass observed: 1454.09 ± 0.18. Charge states; 485.69 [M+3H]^3+^, 727.98 [M+2H]^2+^.

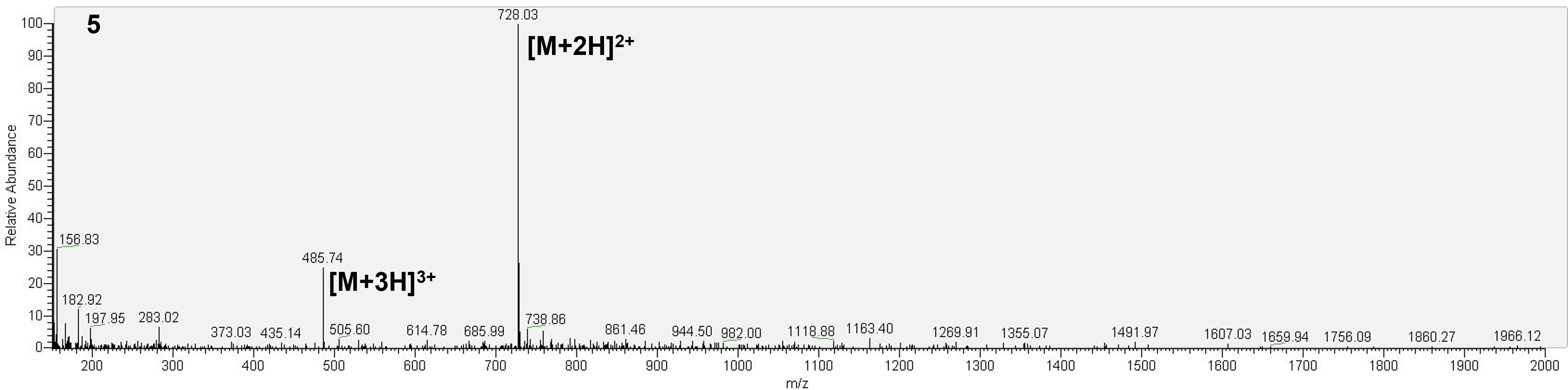

**Supporting Information Figure S28.** ESI-MS of purified peptide, **5**, mass calculated for [C_58_H_83_N_23_O_18_S_2_ + H] 1454.57; deconvoluted mass observed: 1454.14 ± 0.11. Charge states; 485.74 [M+3H]^3+^, 728.03 [M+2H]^2+^.

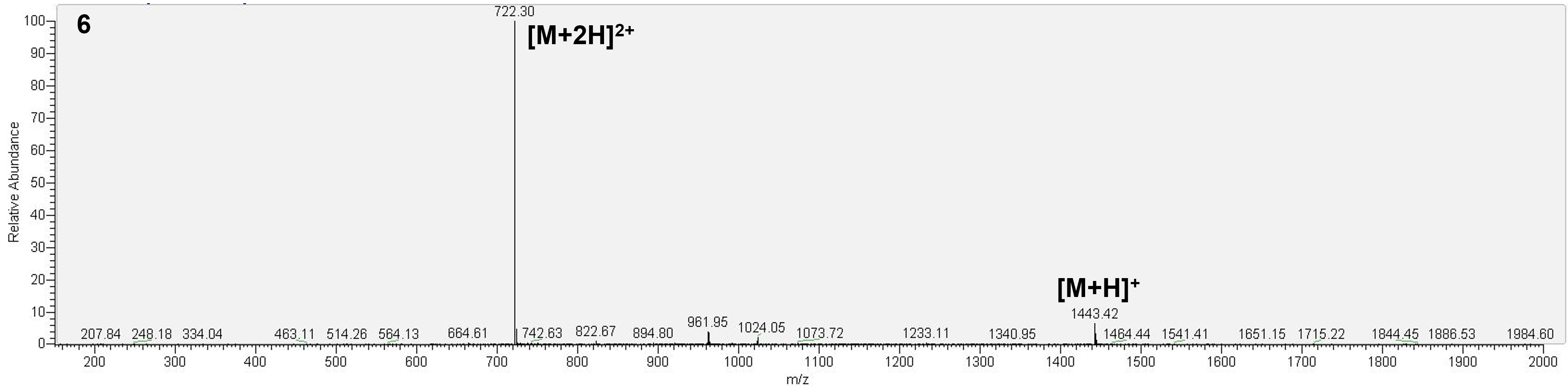

**Supporting Information Figure S29.** ESI-MS of purified peptide, **6**, mass calculated for [C_55_H_83_N_19_O_19_S_4_ + H] 1442.62; deconvoluted mass observed: 1442.51 ± 0.13. Charge states; 722.30 [M+2H]^2+^, 1443.42 [M+H] ^+^.

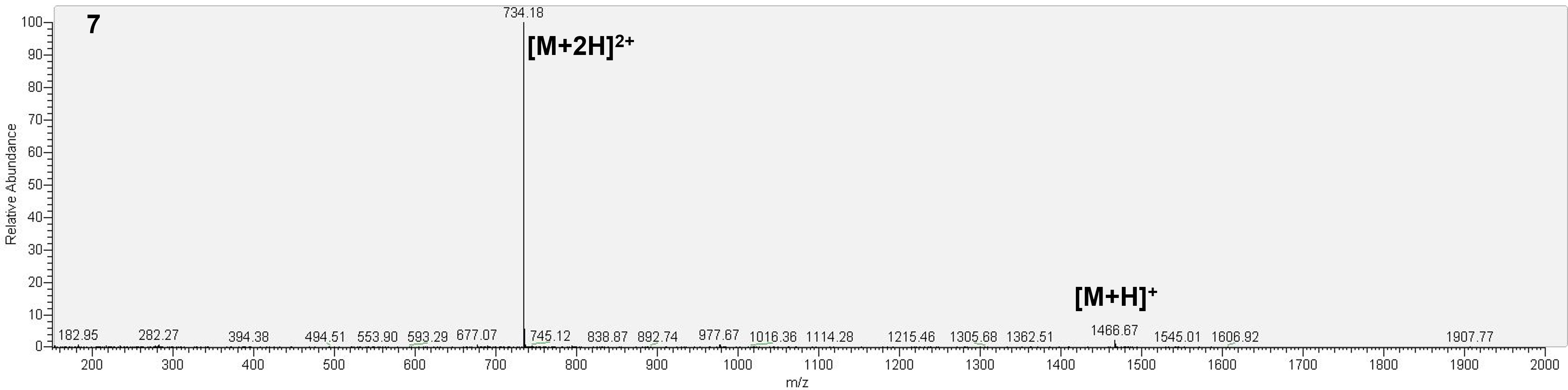

**Supporting Information Figure S30.** ESI-MS of purified peptide, **7**, mass calculated for [C_57_H_83_N_19_O_19_S_4_ + H] 1466.65; deconvoluted mass observed: 1466.02 ± 0.49. Charge states; 734.18 [M+2H]^2+^, 1466.67 [M+H] ^+^.

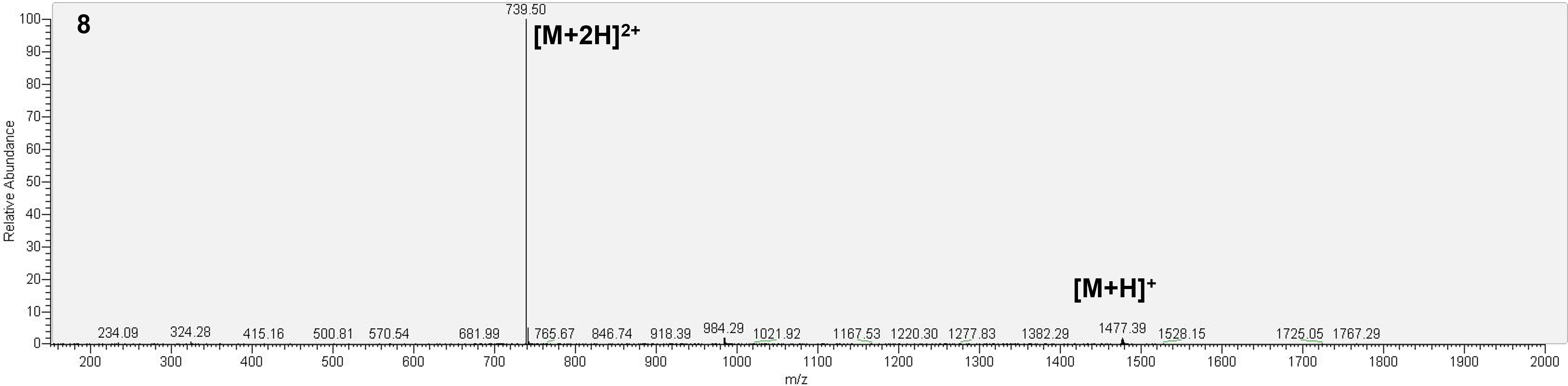

**Supporting Information Figure S31.** ESI-MS of purified peptide, **8**, mass calculated for [C_55_H_82_ClN_19_O_19_S_4_ + H] 1477.07; deconvoluted mass observed: 1476.70 ± 0.43. Charge states; 739.50 [M+2H]^2+^, 1477.39 [M+H] ^+^.

#### Cryo-EM

**Supporting Information Table S3.** Table of cryo-EM data collections and model building statistics (PDBID: 28WS).

|  | **nAChR α-GI:triazole (4)** |
| --- | --- |
| **Data collection** |  |
| EM facility | eBIC |
| Magnification | 165,000x |
| Voltage (kV) | 300 |
| Number of frames | 60 |
| Electron exposure (e^-^/Å^2^) | 60.15 |
| Target defocus (μm) | -0.5 *to* -2.0 |
| Pixel size (Å) | 0.508 |
| Micrographs | 21848 |
| **Reconstruction** |  |
| Initial particles | 851,472 |
| Final particles | 118,387 |
| Symmetry | C1 |
| Box size (pixels) | 512 |
| FSC threshold | 0.143 |
| Map resolution (Å) | 2.66 |
| Map sharpening B factor (Å^2^) | 92.2 |
| **Refinement** |  |
| Overall Q-score | 0.71 |
| Molprobity score | 1.22 |
| Clash score | 4.15 |
| Poor rotamers (%) | 0.00 |
| RMSD values |  |
| Bond lengths (Å) | 0.003 |
| Bond angles (˚) | 0.479 |
| Ramachandran |  |
| Favoured (%) | 97.93 |
| Outliers (%) | 2.07 |
| B-factors (Å^2^) |  |
| Protein | 39.74 |
| Ligands | 73.77 |
| Model composition |  |
| Non-hydrogen atoms | 35526 |
| Protein residues | 2,049 |
| Ligands | 53 |
| Waters | 376 |

**Supporting Information Table S4.** Measured surface area and interfaces of ɑ-GI (**1**) and ɑ-GI mimetic (**4**) at each of the acetylcholine binding sites. A comparison to well-studied toxin ɑ-bungarotoxin is included (PDBID: 28WS).

|  | **Surface Area** | **α_δ_ Interface** | **δ Interface** | **Total Interface** | **Surface Area** | **α_γ_ Interface** | **γ Interface** | **Total Interface** |
| --- | --- | --- | --- | --- | --- | --- | --- | --- |
| **⍺-GI** | 1312.5 | 375.7 | 427.6 | 803.3  (61.2%) | 1328.9 | 372.1 | 447.8 | 819.9  (61.7%) |
| **⍺-GI Mimetic (4)** | 1326.6 | 379.9 | 495.9 | 875.8  (66.0%) | 1359.4 | 407.0 | 487.3 | 894.3  (65.8%) |
| Interface areas are measured in Å^2^. α-GI taken from PDB: 9SRL | | | | | | | | |
